# Spatial selection and local adaptation jointly shape life-history evolution during range expansion

**DOI:** 10.1101/031922

**Authors:** Katrien Van Petegem, J. Boeye, R. Stoks, D. Bonte

## Abstract

In the context of climate change and species invasions, range shifts increasingly gain attention because the rates at which they occur in the Anthropocene induce fast shifts in biological assemblages. During such range shifts, species experience multiple selection pressures. Especially for poleward expansions, a straightforward interpretation of the observed evolutionary dynamics is hampered because of the joint action of evolutionary processes related to spatial selection and to adaptation towards local climatic conditions. To disentangle the effects of these two processes, we integrated stochastic modeling and empirical approaches, using the spider mite *Tetranychus urticae* as a model species. We demonstrate considerable latitudinal quantitative genetic divergence in life-history traits in *T. urticae*, that was shaped by both spatial selection and local adaptation. The former mainly affected dispersal behavior, while development was mainly shaped by adaptation to the local climate. Divergence in life-history traits in species shifting their range poleward can consequently be jointly determined by fast local adaptation to the environmental gradient and contemporary evolutionary dynamics resulting from spatial selection. The integration of modeling with common garden experiments provides a powerful tool to study the contribution of these two evolutionary processes on life-history evolution during range expansion.

## Introduction

Species ranges have always been dynamic. They shrink and expand in response to changing environmental conditions and, accordingly, numerous species are currently shifting their ranges due to contemporary climate change (Parmesan 2006). Moreover, a growing number of species are currently expanding their ranges after being introduced in a new environment by humans (*i.e*. invasive alien species) (Richardson and Rejmanek 2011). During such range expansions or shifts, species undergo multiple selective pressures (Phillips et al. 2010). Especially for poleward range expansions or shifts, a straightforward interpretation of the observed evolutionary dynamics is hampered due to the consequences of both the changing local environmental conditions and the expansion process *per se*. As the feedback between ecology and evolution can occur extremely rapidly (Kubisch et al. 2014), local adaptation and range expansion might occur at similar contemporary timescales, both shaping quantitative genetic trait divergence.

Species expanding or shifting their range poleward experience a change in temperature and growing season, which will affect their life histories. Invasive species, like for example the Colorado potato beetle, change their physiology and diapause behavior in order to cope with the lowered temperatures in more northern regions (Piiroinen et al. 2011; Lehmann et al. 2014; Lehmann et al. 2015). Changes in the length of the growing season, on the other hand, will affect both invasive and niche-tracking species, primarily acting on traits like development time, growth rate and adult size. This is especially true for species with short generation times, having more than one generation per growing season and a cessation of reproduction during winter. The length of the growing season limits the number of generations they can produce per year (*i.e*. voltinism). More specifically, the gradual shortening of the growing season with latitude results in abrupt changes in voltinism and creates a typical saw-tooth pattern in development time (Roff 1980) (see figure 1 for a schematic explanation of this pattern). Since development time is suggested to share an underlying mechanism with growth rate (Kivela et al. 2011), high latitude populations not only tend to compensate for the shorter growing seasons through these changes in development, but also through the evolution of genetically faster growth rates (*i.e*. countergradient variation, see Conover and Schultz 1995). Moreover, development time and growth rate together determine adult size, leading to either bigger, smaller or equal-sized individuals in more northern regions (Blanckenhorn and Demont 2004). Apart from these climatic changes, many range-shifting species may also suffer from changes in habitat quality and quantity. However, this is mainly restricted to native range climate-tracking species (as opposed to invasive species), for which deteriorating habitat is one of the main explanations for the occurrence of their initial range limits (North et al. 2011; Hargreaves et al. 2014).

**Figure 1:**
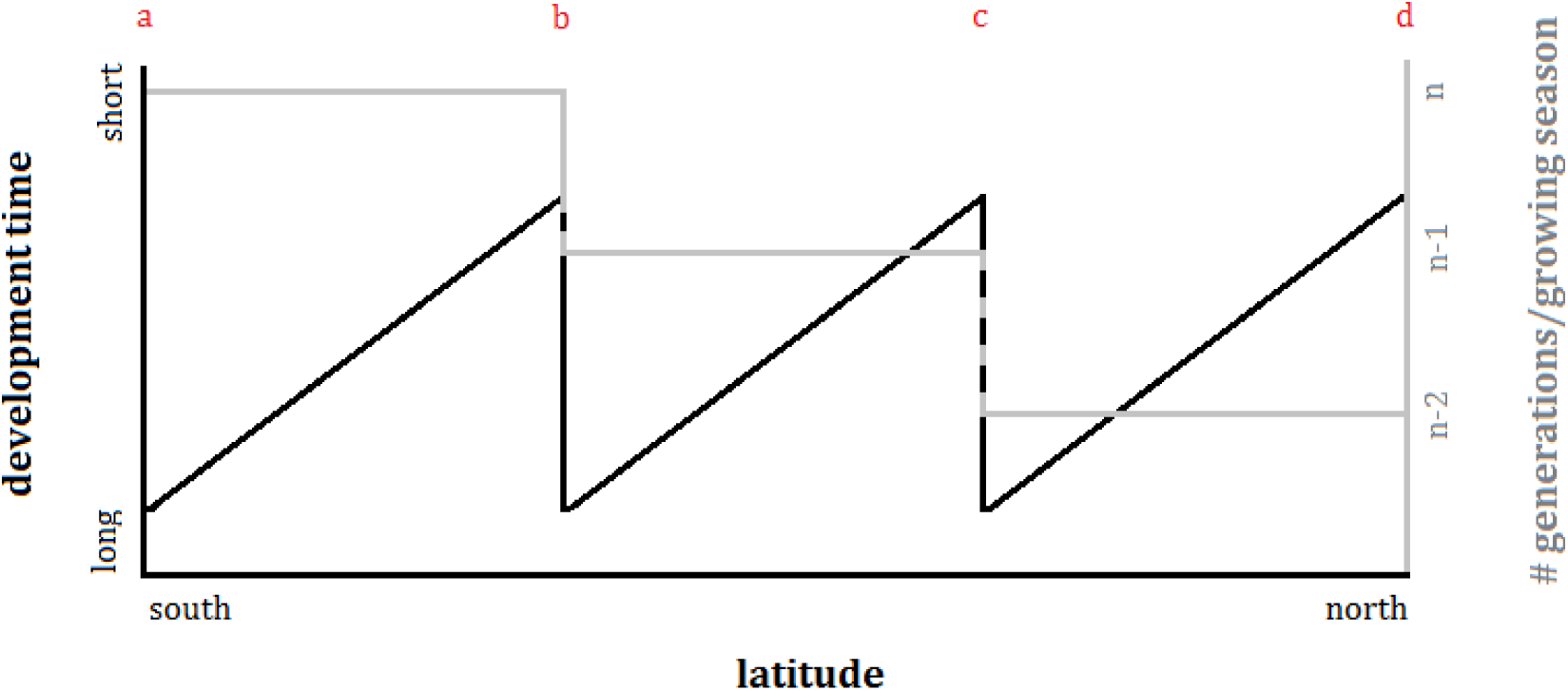
A theoretical scheme of a saw-tooth pattern of development time against latitude for a certain species. At point a, the species has n generations within the growing season (grey line) and its development time is relatively long (slow development) (black line). From point a to b, the number of generations within the growing season remains constant, but the development time of the species increasingly shortens (faster development) because an equal number of generations needs to be produced within an increasingly shorter growing season (shorter growing season towards the north). At point b, the species cannot further speed up its development and 'sacrifices' a generation (shift from n to n-1 generations). Because of this sacrifice, there is a sharp relaxation in development (sudden elongation of development time). From point b to c, the number of generations remains n-1, but the development time increasingly shortens again. At point c, another generation is sacrificed, and from point c to d, development time increases once more.

On top of this pressure to adjust to the changing local environment, the process of range expansion in itself entails a strong selection pressure. Firstly, since the most dispersive phenotypes inevitably accumulate at the expansion front, assortative mating takes place (Phillips et al. 2010; Shine et al. 2011). This results in increased dispersal abilities at the range front, as has already been illustrated theoretically (*e.g*. Travis and Dytham 2002; Burton et al. 2010; Perkins et al. 2013) as well as empirically through field and common garden observations (*e.g*. Phillips et al. 2006; Mitikka and Hanski 2010; Hill et al. 2011; Huang et al. 2015) and experimental evolution (Fronhofer and Altermatt 2015). It is thus acknowledged that dispersal evolution both affects (Kubisch et al. 2014) and is affected by range expansion (reviewed in Hill et al. 2011). Secondly, because of the locally low densities at the leading edge, individuals in the vanguard of an expanding range are predicted to experience *r-* rather than *K*-selection, which would translate into a higher investment in reproductive traits (Phillips 2009; Phillips et al. 2010). Range expansion thus results in a positive selection for dispersal due to the interaction between spatial sorting (and thus assortative mating) of dispersive phenotypes and an increased population growth rate driven by density release at the expansion front. Through both these mechanisms, range expansion therefore contributes to a process of spatial selection (Shine et al. 2011; Perkins et al. 2013).

Local adaptation to a changing climatic gradient and spatial selection thus jointly impact evolutionary dynamics in species expanding poleward. We, however, lack a clear understanding of their relative importance in shaping quantitative genetic trait differentiation along latitudinal gradients. For example, a higher investment in thorax mass in northern populations of an insect species, can result from spatial selection (dispersiveness is selected for at the range front) as well as from local adaptation (lower temperatures might decrease muscle efficiency). Likewise, increased selfing to avoid inbreeding in plants could result purely from local adaptation to ephemeral conditions near the range edge (Hargreaves and Eckert 2014), but could equally be selected for in populations that experience low densities at the expansion front. Insights are, to date, merely derived from theory (Perkins et al. 2013; Hargreaves et al. 2015) or from correlative, often phenotypic, approaches (Therry et al. 2014a; Therry et al. 2014b; Therry et al. 2014c; Therry et al. 2015).

Here, we combine common-garden breeding and an individual-based model to study the causes of multivariate trait evolution during poleward range expansion. Within a full life-history perspective, we assess latitudinal quantitative genetic trait differentiation in the two-spotted spider mite *Tetranychus urticae* Koch (Acari, Tetranychidae), which has recently expanded its European range from the Mediterranean (Carbonnelle et al. 2007) to (at least) southern Scandinavia (personal observation). By contrasting observed empirical patterns in life-history trait divergence with those derived from a stochastic, individual-based simulation model, we are able to assess which evolutionary processes shape the quantitative genetic trait divergence and which specific life history traits are selected by either spatial selection, local adaptation, or their joined action.

## Materials and Methods

### Life history evolution along the sampled gradient

#### Study species

The herbivorous spider-mite *T. urticae* is an agricultural pest species with a worldwide distribution. It reproduces through arrhenotokous parthenogenesis, whereby unfertilized eggs develop into males and fertilized eggs into females. Sex ratio in *T. urticae* is usually female biased (3:1) (Krainacker and Carey 1989), but mothers can alter the sex ratio of their young (Young et al. 1986). Each female may produce more than 50 female offspring, and at optimal temperatures (27-30°C), mites can complete their life cycle in about 8 to 10 days (Sabelis 1981). This contributes to high population growth rates. When foraging, the species engages in short distance ambulatory movements, but it has the ability to disperse long distances (making use of aerial currents) when its food source becomes depleted. Like many arthropods, *T. urticae* can go into diapause when conditions are suboptimal (*e.g*. food shortage, desiccation or cold). This ability is restricted to the adult stage of the species. Over the last decades, the mite species has expanded its European range from the Mediterranean to (at least) southern Scandinavia (personal observation) (more information is provided in Carbonnelle et al. 2007).

#### Population sampling

We collected spider mites during the summers of 2011 and 2012. Based on satellite images, we first selected several collection sites along an 800km latitudinal gradient from north-western Belgium to northern Denmark. In order to minimize variation due to adaptation to different host plant species and human pressure (*e.g*. harvesting, pesticides) and to maximize latitudinal, climatic variation relative to variation in continentality (*i.e*. longitudinal variation), all sampling sites were situated in (semi-)natural area along the coast. All selected sites were visited and mites were searched for on infested leaves of *Lonicera periclymenum* (European honeysuckle), *Euonymus europaeus* (European spindle) and *Humulus lupulus* (common hop), with *L. periclymenum* being a dominant plant species in the north, and *E. europaeus* and *H. lupulus* typically being present in the south (see supplementary material S.1). In 2011, spider mites were found and sampled in twenty different sites (figure 2-more information is provided in supplementary material S.1). In 2012, spider mites were collected in twelve out of these twenty sites, thereby omitting populations that were very close to one another (see supplementary material S.1). To avoid mites being in common garden conditions too long (allowing domestication), trait assessments were split up over two consecutive years. Diapause incidence, longevity, fecundity, egg survival, juvenile survival and development time were assessed with mites collected in 2011, while dispersal propensity, dispersal latency, sex ratio and adult size were assessed with mites collected in 2012.

**Figure 2:**
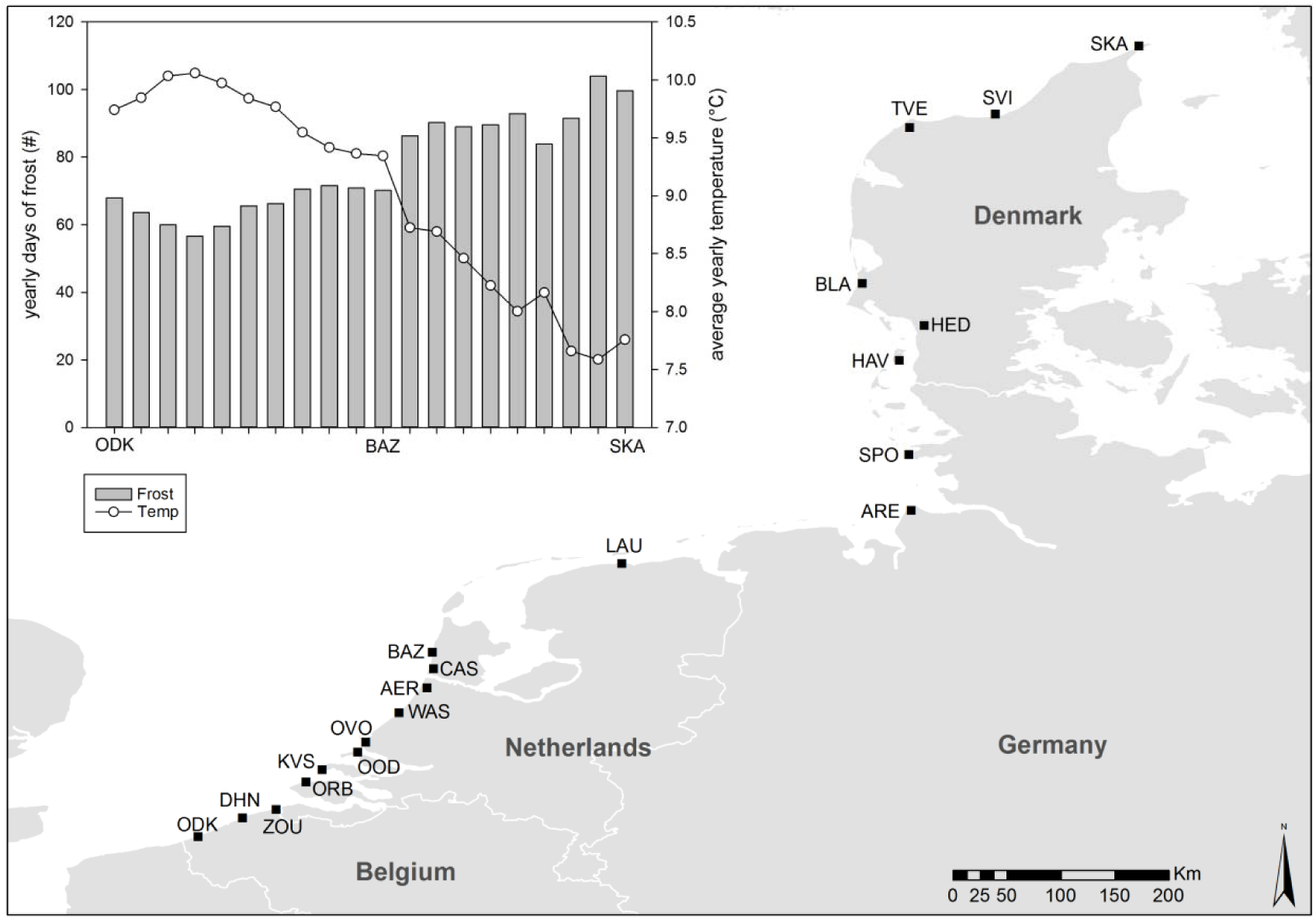
A map showing all the field collection sites in Belgium, The Netherlands, Germany and Denmark. The graph shows the yearly number of frost days and the average yearly temperature for each collection site along the latitudinal gradient. These climatic data were obtained from FetchClimate (Microsoft Research, Cambridge) and were averaged over a period of 35 years (1980 to 2015). (More information is provided in supplementary material S.1.)

#### Common garden and synchronization

Once in the lab, a common garden stock population was each year (*i.e*. in 2011 and 2012) generated for each collection site by putting between fifty and several hundred mites from the collection site on whole bean plants (*Phaseolus vulgaris*, variety Prélude). These common garden stock populations were then maintained at room temperature with a light - regime of 16:8 LD. Mites remained in these stock populations for one to four generations (with the exception of the assessment of sex ratio, where they were in common garden for about 20 generations). Furthermore, before the start of an experiment, a synchronization of the mites was performed to obtain a large pool of same-aged, mated adult females. For this purpose, more than twenty adult females were randomly taken from the stock populations and put on a bean leaf. They were subsequently allowed to lay eggs for 24 hours in a climate room at 27°C, with a light-regime of 16:8 LD. These same-aged cohorts of eggs were then left to develop, and the resulting synchronized adult females were used for all assessments of trait variation. With the exception of diapause incidence (which was performed almost immediately after population sampling), all these assessments were thus performed with at least third generation mites (at least one generation in the stock population, plus one generation of synchronization). Direct environmental effects (including host plant mediated effects and environmentally induced maternal effects) could therefore be excluded.

#### Data collection and statistics

A detailed overview of the applied methodology during data collection can be found in supplementary material S.2. In short, we measured the following ten life-history traits for all the sampled populations: dispersal propensity, dispersal latency, diapause incidence, fecundity, longevity, adult size, egg survival, juvenile survival, sex-specific development time and sex ratio. The first six traits were only assessed for female mites (not relevant for males). Both dispersal propensity and dispersal latency (*i.e*. the period of‘decision-making’ before dispersal) were assessed by means of established behavioral assays (see Li and Margolies 1994; Van Petegem et al. 2015). Dispersal is density-dependent (Harrison 1980; Denno and Peterson 1995) and the density used in our setup could thus affect the outcome. We therefore measured the mites’ propensity to disperse at different densities and tested whether the reaction norm differed between latitudes. We furthermore obtained a population-level measure for intrinsic growth rate by multiplying values of lifetime fecundity with juvenile survival, egg survival and 1/sex ratio (*i.e*. ratio of daughters to total number of offspring). These values were resampled from the quantified distributions of the respective traits. The number of samples used for this purpose was set to the sample size of sex ratio (*i.e*. the lowest sample size of all traits).

Prior to the univariate analyses of the data, we performed a multivariate distance-based ANOVA to test for variation in multivariate life-history parameter space (all ten measured life-history traits), using the vegan and permute packages of R version 3.1.0 (The R Foundation for Statistical Computing, 2014) (see supplementary material S.2 for more detailed information). As this multivariate ANOVA showed considerable variation in life-history strategies among the different sampled populations (F_1,8_=2.2285; p=0.051 for the subset of 12 populations sampled in both 2011 and 2012, and hence all nine measured traits; and F_1,16_=3.6568; p=0.009 for eighteen of the twenty populations sampled in 2011 and hence the subset of six traits measured for only 2011), we then used (generalized) linear mixed models (SAS 9.4, SAS Institute Inc. 2013) to assess, for each trait separately, differentiation along the latitudinal gradient. Latitude, mite density (for dispersal propensity) and host plant species were the independent variables. We tested trait differentiation related to host plant identity because the dominant host plant species in the field changed with latitude and could thus have affected our latitudinal signal. Maternal line was added as a random effect. We again refer to supplementary material S.2 for a detailed outline of the different models. Additionally, to infer the presence of the theoretically predicted saw-tooth pattern for development time, a fourth order polynomial model (*i.e*. shape of saw-tooth pattern) was tested for its fit with the data on development (see supplementary material S.2 for more information).

### Inferring mechanisms by contrasting the empirical data with a parameterized individual-based model

#### The individual-based model

We here only outline the basic principles of our individual-based model, but provide a detailed description and motivation in supplementary material S.3. We designed a stochastic, individual-based and spatially explicit model to simulate the evolutionary dynamics in *T. urticae* along a climatic gradient comparable to the one we studied empirically. Our model is inspired by that of Bancroft and Margolies (1999), who used an individual-based model to simulate the dynamics among *T. urticae*, its host plant and its predator. We adjusted this existing model to simulate population dynamics at a time-step basis of one day (for which empirical data were available), in a simplified model landscape. This landscape consisted of one hundred rows (latitude) and five columns (longitude) of 10x10 km^2^ grid cells, and population dynamics were consequently scaled to this spatial dimension. This simulated latitudinal range of 1000 km roughly corresponds with the macro-geographic scale at which we sampled *T. urticae* in the field (see earlier). The length of the growing season, determined by seasonal variation in temperature, was defined at the grid-level by two trigonometric functions, which were made using actual data.

Our model simulates the behavior and life history of adult female mites, as females are the reproducing sex and adult females the main dispersers. No mating limitations and recombinations were consequently implemented. Using data from Sabelis (1981), development, longevity, mortality, and fecundity of the mites were all simulated according to the local grid cell temperature. Mites followed a pattern of exponential growth. However, as a compromise to keep computational efficiency, individuals were randomly deleted as soon as more than twohundred mites occupied a grid cell. We neglected short-distance movements and assumed density-independent aerial dispersal, as the large spatial scale used (*i.e*. 10×10km^2^ grid cells) did not allow to sensibly incorporate density-dependence (which is important at the level of a single leaf or plant). We assured rare long distance dispersal by setting a relatively low dispersal mortality (90%, see De Roissart et al. 2015). The probability for an individual mite to engage in arial long-distance dispersal was modeled as an unconditonal nearest-neighbour dispersal rate, determined by a single locus subject to selection/mutation. Other traits subject to selection and mutation were development, fecundity and the timing of diapause onset and termination. A linear trade-off between development and fecundity was implemented in order to constrain the evolutionary trajectories. This trade-off function was coded by a single-locus trait that altered the balance of investment between fecundity and development, by ‘taking resources’ from one trait and ‘investing them’ in the other. The maximal increase or decrease in performance of either trait was limited to 10 or 20% (assumed realistic, conservative percentages). We tested all possible combinations of these trade-off limitations because no empirical data on this kind of trade-off are available.

Because we aimed to contrast evolutionary dynamics resulting from spatial selection and local adaptation, we tested three competing scenarios: (1) a scenario of range expansion along a homogeneous gradient, (2) a scenario with range expansion along a latitudinal climatic gradient and (3) a scenario where evolution occurs within this same heterogeneous gradient but without the process of range expansion. In the stable range scenario (scenario 3), individuals were initialised along the entire climatic gradient. For scenarios with evolution during range expansion (senarios 1 and 2), only the ten ‘southernmost’ rows were initialised with genetically diverse individuals, thereby allowing range expansion towards the ‘northern’ grid cells. This range expansion was constrained in scenario 2 by the seasonal conditions that affect development, survival and fecundity (from Sabelis 1981; see supplementary material S3). Evolutionary dynamcis were in equilibrium after 100,000 time-steps (days), corresponing with a total time of 273 years. The entire range was occupied after approximately 30,000 time steps in the two range expansion scenarios.

#### Comparison of empirical and simulation results

While our developed individual-based model is one of the most detailed to date that allows the prediction of eco-evolutionary dynamics in a model species, it lacks mechanistic details regarding the mode of inheritance, scaling, trait responses to common garden conditions and impacting local environmental pressures (other than climate). We therefore followed a pattern-oriented approach (Grimm et al. 2005), by means of Approximate Bayesian Computation (Csillery et al. 2010; Baiser et al. 2013; Wiegand and Moloney 2014), that allowed us to compare emerging statistical properties of the three contrasting model scenarios (see above) with those empirically derived. Because we expected the outcome of our approach to depend on the implemented trade-off function between fecundity and development time (see above and supplementary material S.3, S.3.5), we also included an analysis for this trade-off. We therefore varied the maximal costs and benefits for development time and fecundity in all combinations of 10% and 20% (10%-10%, 10%-20%, 20%-10%, 20%-20% *-e.g*. 10%-20% implies that a *max*. 10% increase/decrease in development corresponds with a *max*. 20% decrease/increase in fecundity), and tested which trade-off setting gave the best fit with our empirical data and was thus most realistic. As such, we compared the goodness-of-fit between our empirical data and each of twelve models: three model scenarios (either a stable range with latitudinal climatic variation, an expanding range in a climatic homogenous environment or a range expansion scenario along the climatic gradient), each with four possible trade-off balances. This goodness-of-fit was derived for statistical patterns of those three traits that were subject to spatial and natural selection and for which a comparison between simulation and empirical data could be made (*i.e*. the regression slope against latitude of intrinsic growth rate, dispersal, and develoment time, and the amplitude and wavelength of the saw-teeth in the saw-tooth pattern in development time; see supplementary material S.4). We randomly sampled one out of 100 simulated values from the simulation enveloppe and calculated its deviance with the empirically observed value (*e.g*. sampling one out of the 100 simulated values for the regression slope in dispersal and calculating the deviance of this value with the empirical value of the regression slope in dispersal). The model scenario with the lowest deviance was considered to have the best goodness-of-fit with the empirical data for the specific statistical pattern. The probability for each model to have the best goodness-of-fit for a specific statistical pattern was achieved by repeating this procedure 10,000 times. Finally, to infer the best-matching evolutionary scenario (range expansion *vs*. stable range and environmental gradient *vs*. no gradient) for each of the statistical patterns, we calculated Bayes’ factors (over all four trade-off settings). A Bayes’ factor of three or more for a model comparison of model A *versus* B, implies that model A is more strongly supported by the data (Kass and Raftery 1995).

## Results

### Life history evolution along the sampled gradient

#### Dispersal propensity and latency

Dispersal propensity and latency were both significantly affected by latitude: dispersal propensity increased with latitude (F_1,2235_=33.93; p<0.0001) (figure 3A), while dispersal latency showed the exact opposite trend (F_1,469_=12.16; p=0.0005) (figure 3B). Dispersal propensity and latency were density-dependent, but this density-dependence was not affected by latitude (propensity: F_2,2230_=0.03; p=0.9702 and latency: F_2,467_= 2.71; p=0.0678). There was no effect of host plant species on dispersal propensity (F_3,2232_=1.85; p=0.1356), nor on dispersal latency (F_3,464_=0.60; p=0.6164).

**Figure 3:**
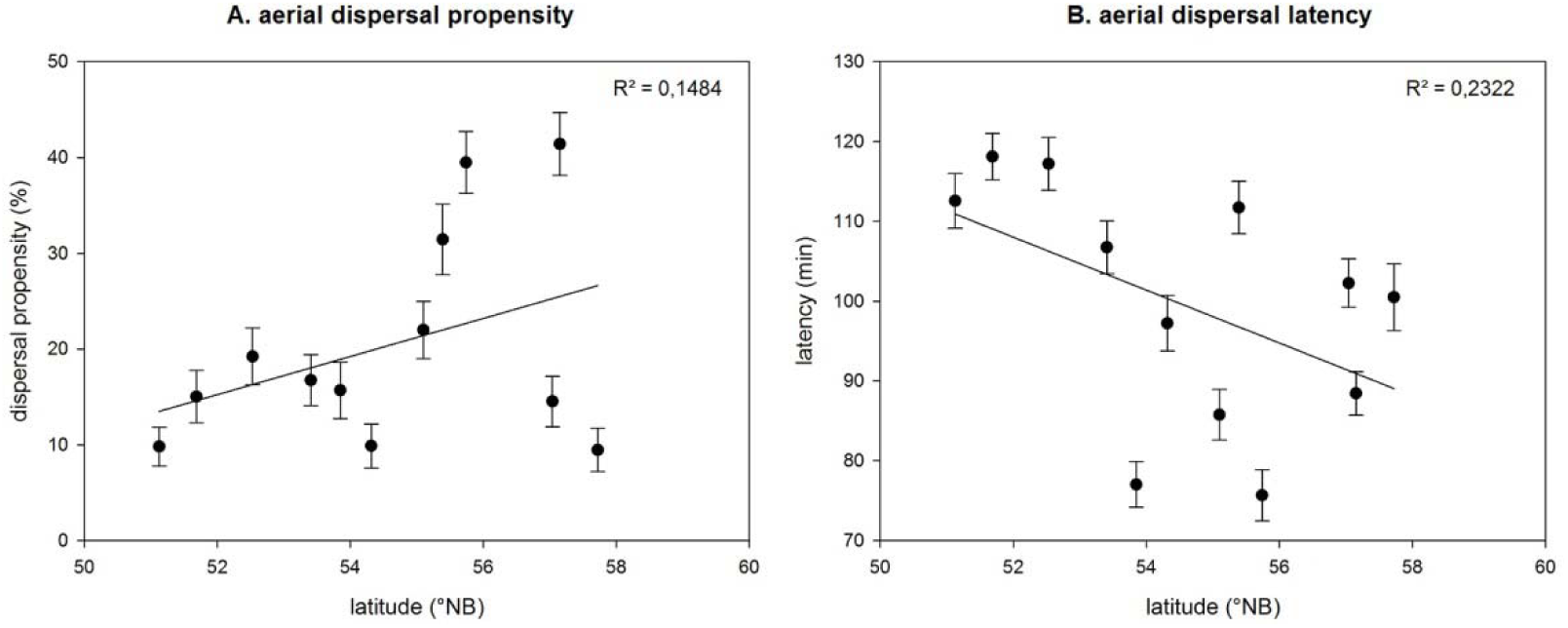
Dispersal propensity (A) and dispersal latency (B) for each sampled population along the latitudinal gradient. Populations means are given ± 1 standard error (bars). Regression lines are shown together with there Revalues. These values are calculated in a conservative way, using the population averages.

#### Diapause incidence

No effects of latitude on diapause incidence were found (F_1,18,68_= 0.05; p=0.8233) (see supplementary material S.5). Instead, the diapause incidence of the mites was significantly affected by the host plant on which the mites were collected (F_3,50.13_=9.86; p<0.0001) (see supplementary material S.5).

#### Fecundity and longevity

Lifetime fecundity (F_1,71_=14.75; p=0.0003) (figure 4A) and longevity (F_1,65,1_=11.41; p=0.0012) (figure 4B) both decreased significantly with an increasing latitude. For daily fecundity, however, no effects of latitude were found (F_1,68_=0.69; p=0.4103). Instead, daily fecundity was affected by the host plant species (F_3,69_=5.59; p=0.0017) (see supplementary material S.5). For lifetime fecundity, no effects of host plant species were found (F_3,68_=1.62; p=0.1932). For longevity, there was a general effect of host plant species (F_3,66,3_ = 3.72; p=0.0155), but none of the adjusted p-values were significant in the pairwise post-hoc Tukey tests.

#### Egg survival, juvenile survival and development time

The relative amount of hatched eggs increased significantly with latitude (F_1,103,1_=6.76; p=0.0107) (figure 4C), but the proportion of juvenile mites reaching the adult life stage showed no latitudinal pattern (F_1,1315_=0.19; p=0.6663) (see supplementary material S.5). Furthermore, towards higher latitudes, female (F_1,66,1_=11.03; p=0.0015) and male (F_1,62,1_=18.84; p<0.0001) spider mites had a significantly shorter development time (*i.e*. a faster development) (figure 4D). There was no effect of host plant species on the development time of females (F_3,57,4_=1.89; p=0.1416) or males (F_3,60_=2.21; p=0.0960), nor on egg survival (F_3,44,38_=2.51; p=0.0706) or juvenile survival (F_3,1312_=1.90; p=0.1277).

The fourth order polynomial fitted the data on development time significantly (F_1,31_=6.68; p=0.0147). It explained 57% (for males) and 27% (for females) of the variance. The significance of the fourth order polynomial indicates that our empirical data followed a saw-tooth pattern (see figure 3D).

#### Sex ratio and adult size

From more southern to more northern latitudes, the sex ratio of the sampled populations increased significantly (F_1,61,97_=6.73; p=0.0118) (figure 4E). With increasing latitude, populations were thus increasingly male-biased. Adult size, in contrast, was not affected by latitude (F_1,342_=1.19; p=0.2761) (see supplementary material S.5). Instead, the adult size of the female spider mites was significantly affected by the host plant species from which they were collected (F_3,343_ = 3.64; p=0.0130) (see supplementary material S.5). There was no effect of host plant species on sex ratio (F_3,50,9_=2.10; p=0.1124).

#### Intrinsic growth rate

Intrinsic growth rate decreased significantly towards higher latitudes (t_9_=-2.49; p=0.0376) (figure 4F).

**Figure 4:**
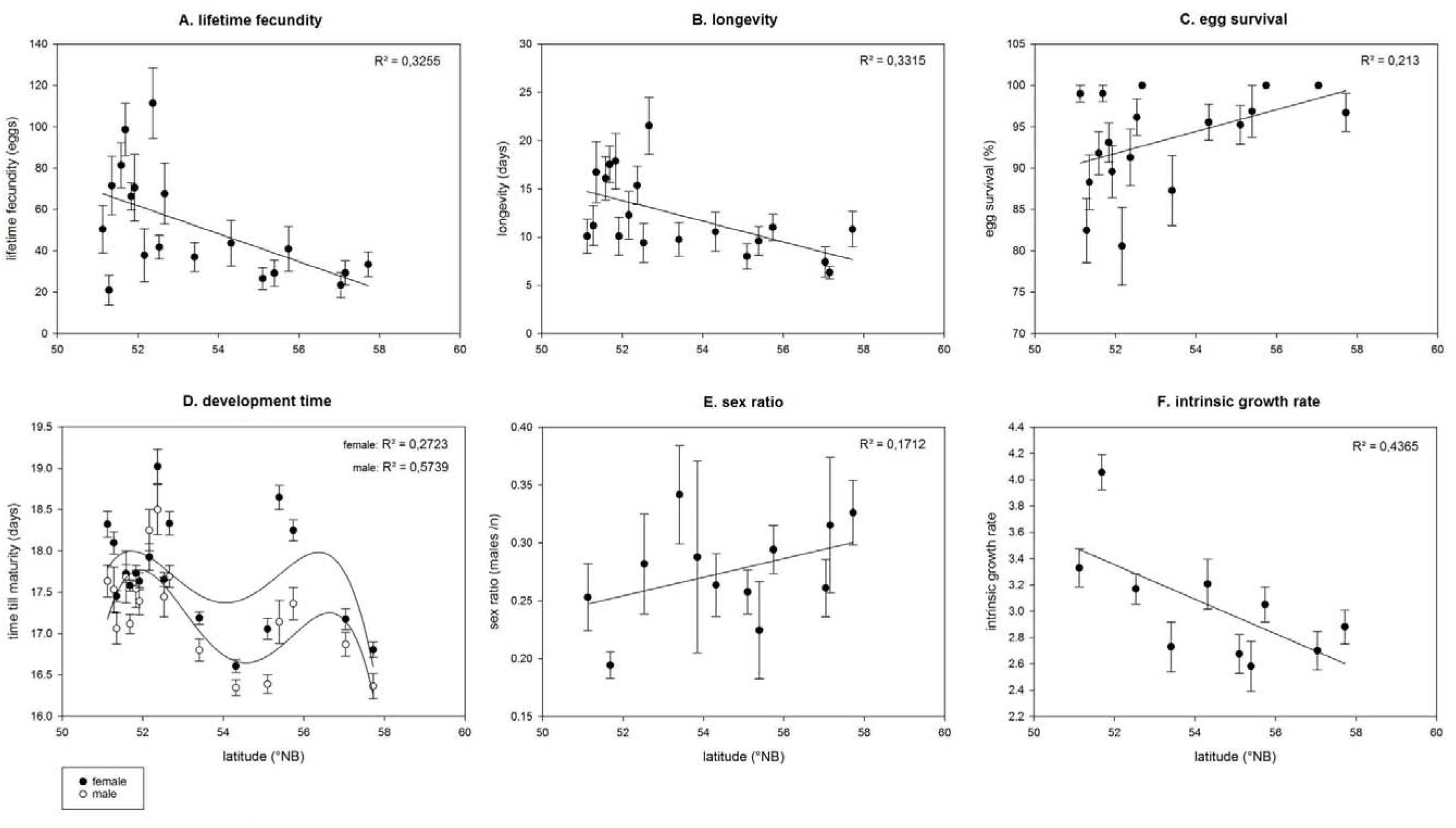
Lifetime fecundity (A), longevity (B), egg survival (C), development time (D), sex ratio (E) and intrinsic growth rate (F) for each sampled population along the latitudinal gradient. Population means are given ± 1 standard error (bars). Regression lines (fourth order polynomial in case of figure 4D) are shown together with there Revalues. These values are calculated in a conservative way, using the population averages. In figure 4D, development time is shown separately for females (closed circles) and males (open circles).

### Inferring mechanisms by contrasting the empirical data with a parameterized individual model

Three consistent (*i.e*. consistent over all the trade-off settings) patterns in life-history divergence along the latitudinal gradient emerged from the model: an increase in dispersal towards the range front in the range expansion scenarios, a stepwise decrease in voltinism towards the north in the scenarios with an environmental gradient and an overall lower temperature for diapause termination than for diapause onset in all scenarios (see supplementary material S.6). Our results furthermore show that the chosen trade-off setting in our model (max. effect on development *vs. max*. effect on fecundity) affected voltinism and the relative investment in development *vs*. fecundity (see supplementary material S.3.5), and as such the goodness-of-fit of our three model scenarios for the statistical patterns in development time and intrinsic growth rate (table 1 and figure 5).

The goodness-of-fit for the five different statistical patterns clearly showed differences between the three competing model scenarios (table 1 and figure 5). The stable range scenario poorly predicted the empirically observed dispersal propensity, but provided the strongest supports for the pattern in development time. Overall, however, the stable range scenario performed rather badly. The scenario with range expansion in a homogeneous landscape showed a moderate overall fit, but provided the strongest support for the empirical pattern in intrinsic growth rate. The scenario with range expansion along an environmental gradient resulted in the best overall fit, with the highest values for dispersal propensity and good to strong support for the patterns in intrinsic growth rate and development time. In terms of evolutionary scenario (range expansion *vs*. stable range and environmental gradient *vs*. no gradient), the range expansion scenarios clearly provided a much stronger support for the empirical pattern in dispersal than the stable range scenario (Bayes’ factor =40.15), while no difference in support was found between the scenarios with and without an environmental gradient (Bayes’ factor =0.79). In contrast, the best support for the empirical pattern in development time was provided by the scenarios with an environmental gradient (especially concerning the slope (Bayes’ factor =1.99) and amplitude (Bayes’ factor =3.66) of the pattern), while no difference in support was found between model scenarios with or without range expansion (slope: Bayes’ factor =0.56, amplitude: Bayes’ factor =0.56). Regarding intrinsic growth rate, no clear difference between the evolutionary scenarios was found (range expansion *vs*. stable range: Bayes’ factor =1.41; gradient vs. no gradient: Bayes’ factor =0.74), though the two best fits with the empirical data were provided by range expansion models (see table 1).

**Table 1:** Results from the goodness-of-fit analysis

| model parameters |  |  |  | fitting analysis results (% best fit) |  |  |  |  |
| --- | --- | --- | --- | --- | --- | --- | --- | --- |
| range<br>dynamics | environmental<br>gradient | *max.* %<br>trade-off effect<br>development | *max.* %<br>trade-off effect<br>fecundity | dispersal<br>propensity<br>(slope) | intrinsic<br>growth rate<br>(slope) | development<br>rate<br>(slope) | development<br>rate<br>(wavelength) | development<br>rate<br>(amplitude) |
| stable | yes | 20 | 10 | 0.29 | 7.71 | 0.54 | 14.38 | 1.32 |
| stable | yes | 20 | 20 | 0.44 | 0.61 | 0 | 7.17 | 29.48 |
| stable | yes | 10 | 10 | 0.3 | 10.59 | 5.92 | 6.57 | 15.41 |
| stable | yes | 10 | 20 | 0.2 | 7.26 | 40.56 | 0.86 | 1.09 |
| expansion | no | 20 | 10 | 11.23 | 25.08 | 3.66 | 7.24 | 0.89 |
| expansion | no | 20 | 20 | 5.29 | 5.66 | 3.81 | 5.66 | 5.81 |
| expansion | no | 10 | 10 | 11.79 | 7.66 | 5.1 | 11.86 | 1.02 |
| expansion | no | 10 | 20 | 10.34 | 1.88 | 7.53 | 6.32 | 4.3 |
| expansion | yes | 20 | 10 | 11.55 | 18.19 | 4.98 | 12.08 | 6.59 |
| expansion | yes | 20 | 20 | 14.24 | 6.38 | 6.22 | 10.69 | 14.85 |
| expansion | yes | 10 | 10 | 15.36 | 4.54 | 6.58 | 10.2 | 5.78 |
| expansion | yes | 10 | 20 | 18.97 | 4.44 | 15.1 | 6.97 | 13.46 |

The results from goodness-of-fit analyses between our empirical data and our three competing model scenarios (each with four possible trade-off settings, resulting in a total of twelve models). At the left side of the table, the model parameters are listed (range dynamics, presence or absence of an environmental gradient and the trade-off settings: all combinations of a *max.* $\pm 10\%$ and a *max.* $\pm 20\%$ trade-off effect on development or fecundity). At the right side, the results of the analyses are shown. The percentages indicate how often a model provided the best fit to a particular empirical statistical pattern. (The five statistical patterns: the regression coefficient of the slope in dispersal propensity, the regression coefficient of the slope in intrinsic growth rate, the regression coefficient of the slope in development rate, the wavelength of the saw-teeth of the saw-tooth pattern in development rate and the amplitude of the saw-teeth of the saw-tooth pattern in development rate).

**Figure 5:**
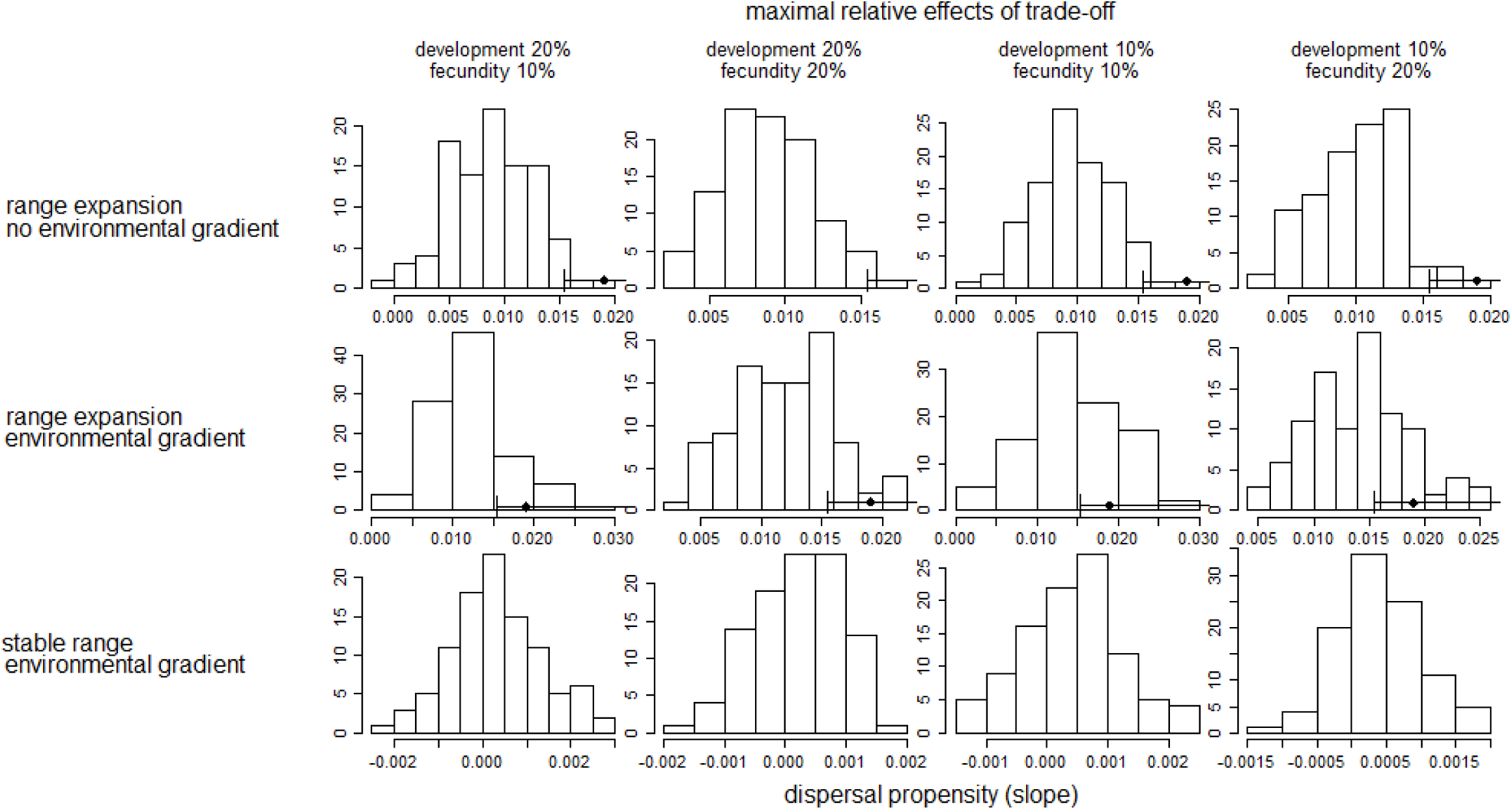
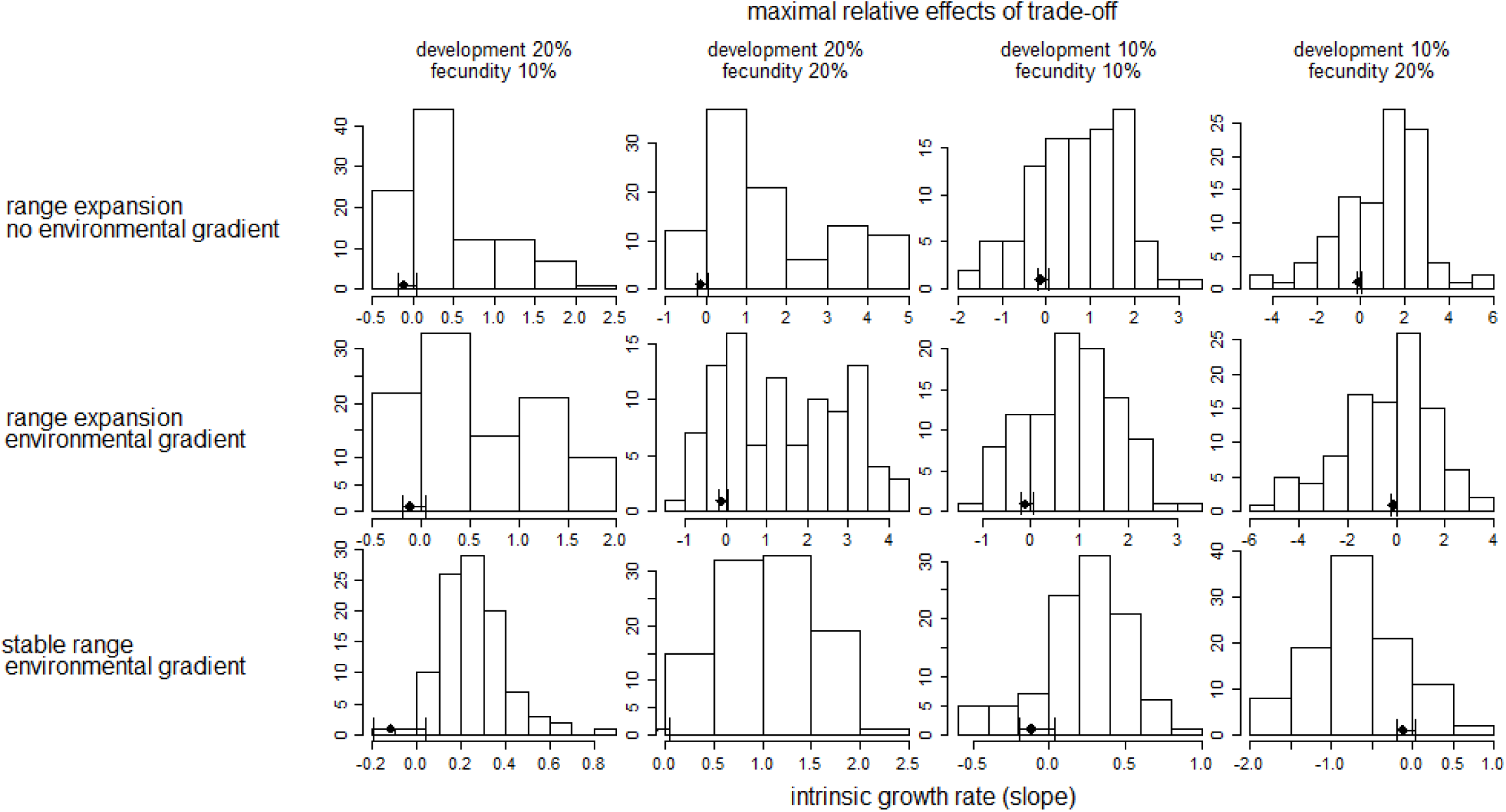
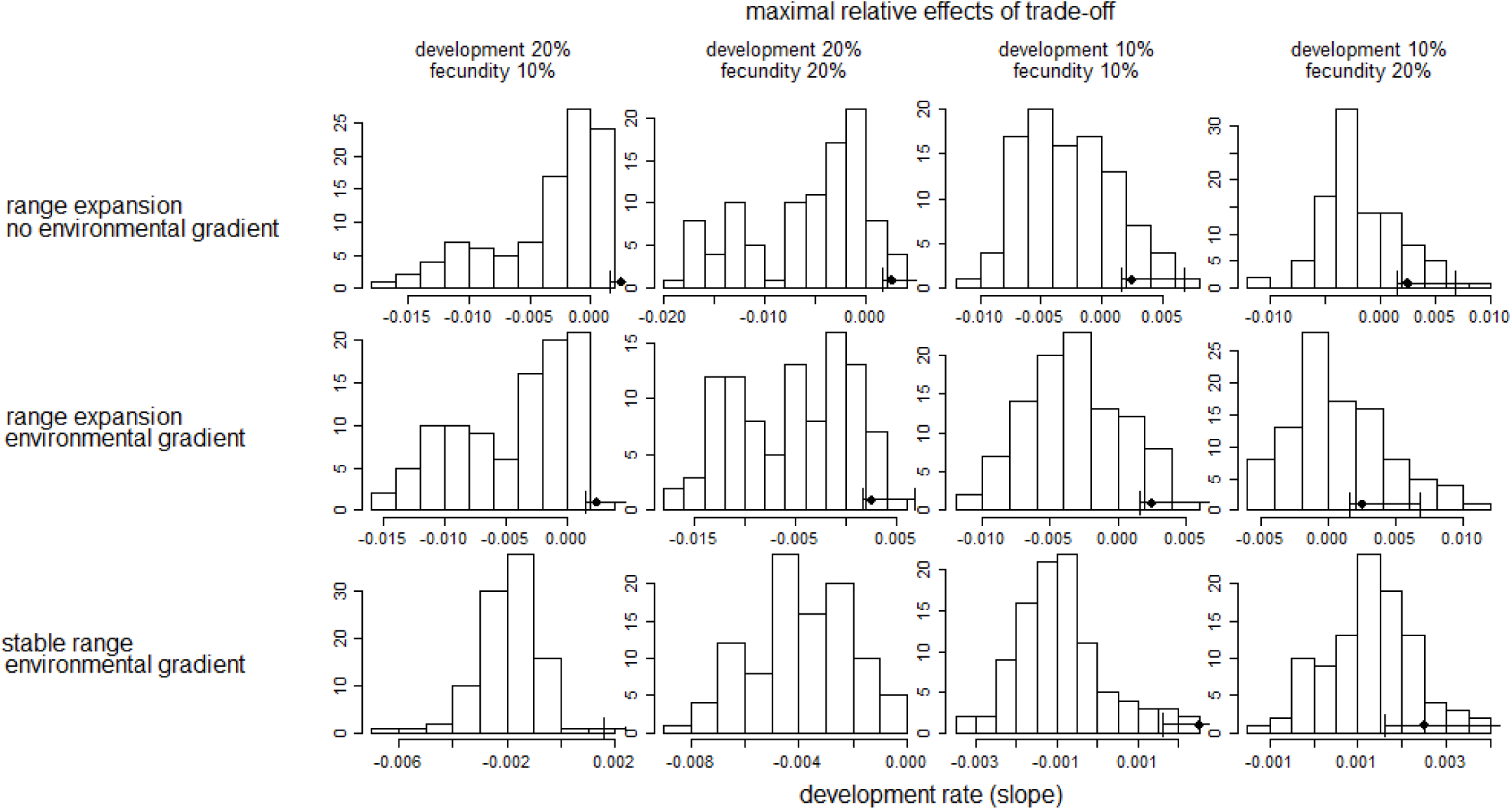
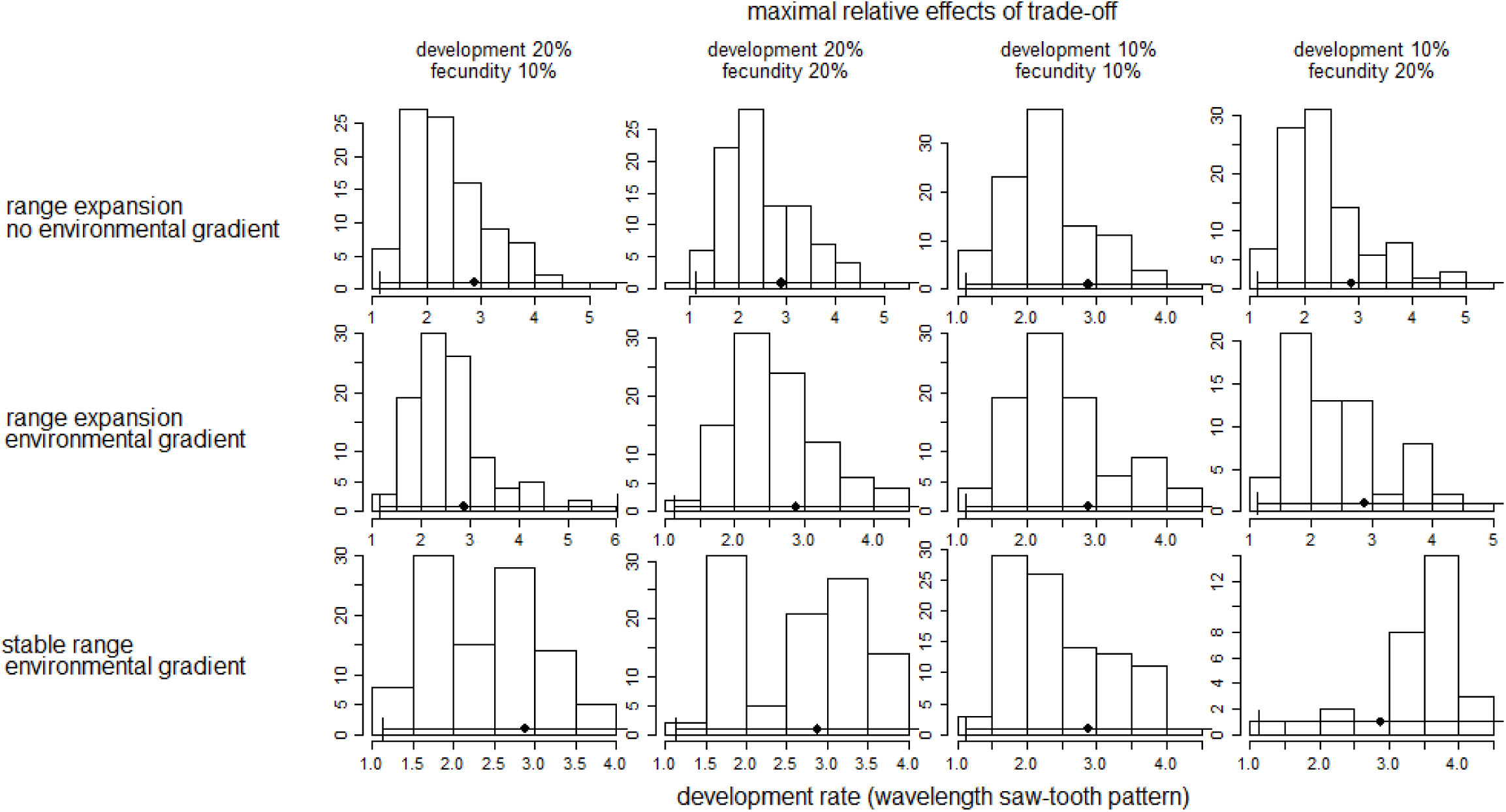
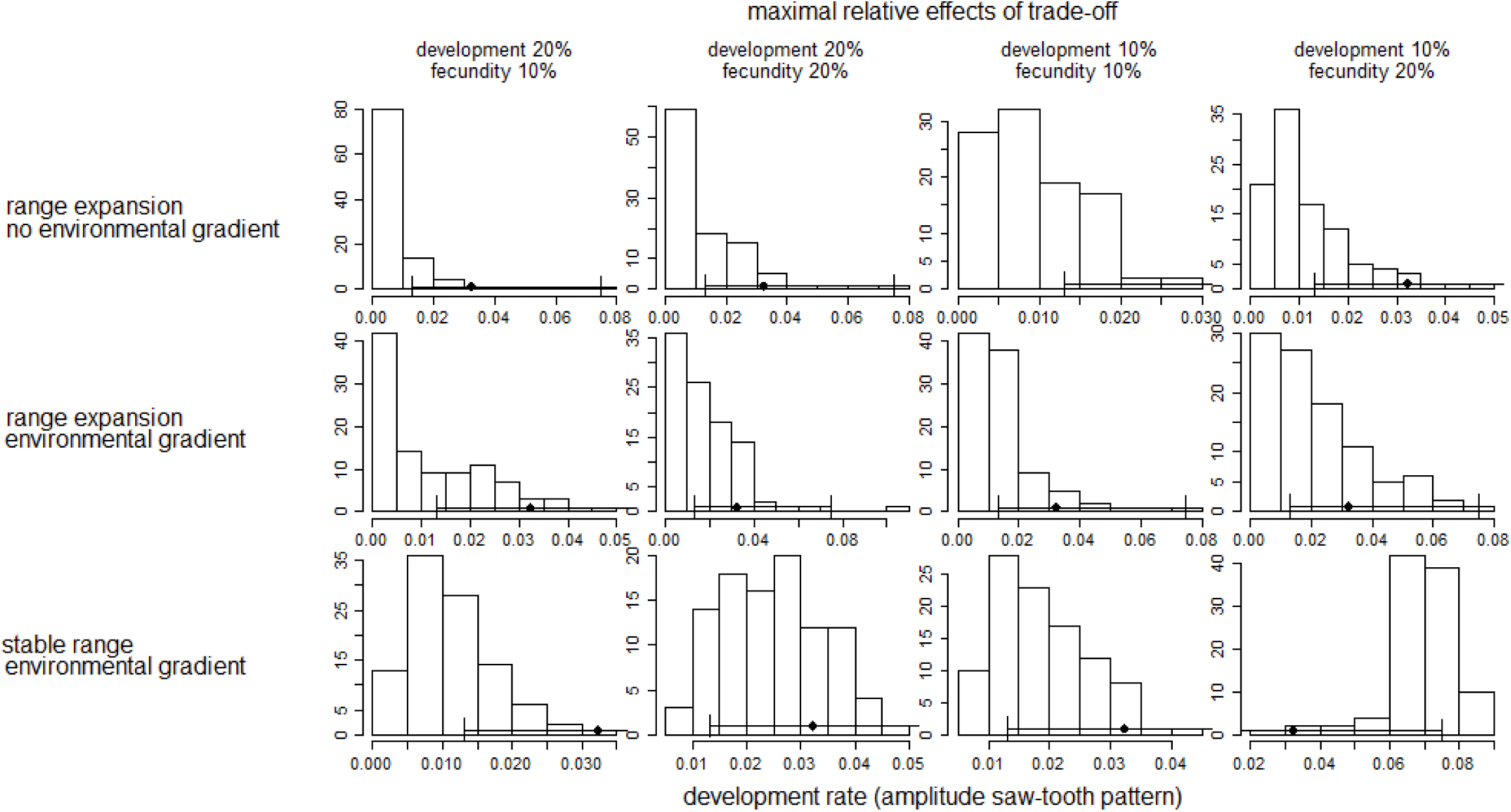
The distribution of 100 simulated results for 5 statistical patterns [the regression slope of dispersal propensity, intrinsic growth rate and development rate, and the wavelength and amplitude of the teeth in the saw-tooth pattern in development rate) in the three model scenarios and for four trade-off settings [10%-10%, 10%-20%, 20%-10% and 20%-20% *max*. relative trade-off effect on development and fecundity, respectively). The black diamond represents the empirical value for the statistical pattern [absence means that the value is outside the distribution of simulated results) and the bars show the 95% credibility interval around this value [calculated through bootstrapping). Slopes are calculated from a linear regression over the latitudinal gradient Wavelength and amplitude are calculated for the saw-teeth of the saw-tooth pattern in development rate [see supplementary material S.4). Note that the x-axes have a different scale on each of the figures.

## Discussion

Our common garden approach revealed considerable quantitative genetic divergence in life-history traits in populations of *T. urticae* that were sampled along a latitudinal gradient from range core to expansion front. Dispersal, sex-ratio, egg survival, fecundity, longevity, development time and the derived intrinsic growth rate showed strong latitudinal patterns. By means of pattern-oriented modeling, we demonstrated that increased dispersal at high latitudes resulted from spatial selection, while latitudinal variation in development time was best explained by adaptation to the local climatic and seasonal conditions. Local adaptation and spatial selection consequently jointly shaped quantitative genetic divergence in life history in a poleward range expanding arthropod.

The empirically observed increased dispersal at the range front is in line with several studies on post-glacial range expansion (Cwynar and Macdonald 1987), invasions (Travis and Dytham 2002; Phillips et al. 2006; Huang et al. 2015) and climate change (Thomas et al. 2001; Travis et al. 2013). As this pattern matched best with our range expansion scenarios, this indicates that dispersal ability is positively selected at the front through the process of spatial selection, and not by local adaptation to local environmental conditions (*i.e*. local temperature and growing season length in our model). Evolution of dispersal along a latitudinal gradient could, however, be equally affected by factors related to changes in habitat quality and connectivity (Bowler and Benton 2005), which were not included in our modeling framework. A decrease in habitat quality and connectivity, however, is theoretically expected to select against dispersal (Moran and Alexander 2014), so this would oppose our findings. An additional factor leading to increased dispersal and colonization rates could be the ephemeral nature of range populations (Duputie and Massol 2013), as for example found in a plant species (Darling et al. 2008). However, in our study, host plants were readily available at the range front and expected to be exhausted more slowly than in the range core due to a lower intrinsic growth rate of the mites (see Figure 4F). Increased dispersal at the range edge could also have been caused by increased temporal dynamics (McPeek and Holt 1992), resulting from harsh climatological conditions, especially during winter. In our model, however, this disturbance (see supplementary material S.3.0) was implemented in both the stable and a range expansion scenario and can therefore not explain the difference in dispersal between these two scenarios. Edge populations may furthermore suffer from Allee effects and increased kin competition. Allee effects, however, would result in decreased rather than increased dispersal rates (Travis and Dytham 2002). Kin competition, in contrast, has already been shown to be an important driving force of range expansion (Kubisch et al. 2013). Its relative importance, however, depends on the environmental conditions and it most probably operates in parallel with the process of spatial selection. Spatial selection can thus be considered as a major driver of the evolution in dispersal in our study.

The empirically found latitudinal variation in development time (slope and saw-tooth statistics) matched best with the scenarios that included adaptation towards an environmental gradient. The gradual shortening in the growing season from core to edge, resulted in changes in voltinism and consequent abrupt changes in development time. Indeed, changes in development time are most effective for maintaining an optimal reproductive outcome when a restricted growing season leads to changes in voltinism (Roff 1980). Such adaptations to local conditions are important. When populations are not locally adapted, their population growth rate could slow down drastically, leading to local extinction (Van Dyck et al. 2015). The changes in development time did not cause changes in adult size in our study. This suggests that compensatory growth maintained a constant size at maturity despite large changes in the length of the growing season (Conover et al. 2009). Interestingly, this might imply an increased foraging efficiency at the range margin and thus contradicts predictions of a dispersal-foraging trade-off, found during experimental evolution in a protist (Fronhofer and Altermatt 2015).

Concerning intrinsic growth rate, none of the three model scenarios gave a markedly better fit. The best fits, however, were provided by model scenarios with range expansion. Our empirically observed trend of decreased fecundities in the north, however, opposes theoretical expectations of evolution towards higher intrinsic growth rates at the expansion front, where on average lower population densities occur (Phillips 2009; Phillips et al. 2010). While Fronhofer and Altermatt (2015) showed that the presumption of lower densities at the range margin should not be universally true, we anticipate this to be the case in our study because of the overall shorter growing season and colder temperatures in the north. We therefore tend to attribute the observed pattern in growth rate to trade-offs with other life history parameters. However, while some studies suggest dispersal to trade off with fecundity (*e.g*. Zera and Denno 1997; Hughes et al. 2003), others have failed to detect this, or even found a positive correlation (*e.g*. Saastamoinen 2007; Therry et al. 2015). Furthermore, diapause incidence has been shown not to constrain changes in fecundity in *T. urticae* (Ito 2009). We conducted our study within a full life-history perspective and tested for population-level correlations between multiple life-history traits but did not detect any relevant trade-off (see supplementary material S.7).

The use of common garden breeding techniques is strong to detect genetic variation in traits. We selected our common garden conditions to be as neutral as possible, but different populations (genotypes) may still have reacted differently towards these standardized conditions and may thus have been prone to genotype-by-lab-environment interactions. Full insights into this genetic variation in plasticity, however, can only be achieved by subjecting all populations to a wide range of environmental conditions. Evidently, this would have been logistically impossible for our model system. We thus minimized this bias in our analyses by following a pattern-oriented approach, thereby avoiding a direct comparison of empirical trait values with modeled ones and by using beans as common garden host plants. Beans are known to be a highly favorable environment for spider mites and mites are therefore not expected to show substantial genetic variation in fitness on this plant species (Agrawal et al. 2002; Gotoh et al. 2004). Concerning temperature, however, the relatively high temperatures in our setup (20°C or 27°C) might have induced a slightly bigger shift in environment for mites from northern compared to southern latitudes. In such a changed environment, a higher proportion of males (the genetic equivalent of haploid recombinant genomes) might have been selected for, because it provides a faster response to selection and thus a more rapid adaptation to the new environment (Hartl 1971; Griffing 1982; Havron et al. 1987). Our setup might thus have triggered northern females to produce a more male-biased offspring, as was visible in our empirical data.

Apart from adaptations in function of the latitudinal climatic gradient, we also retrieved signals of local adaptation of the mites towards the host plant species they were collected on. *T. urticae* is known to adapt to new host plant species within ten to fifteen generations (Magalhães et al. 2007), but we only kept the mites in common garden for two to five generations (except for the assessment of sex ratio). This was not sufficient to disrupt adaptation to the previous host plant species (Magalhães et al. 2011). However, the relatively short stay in our common garden setup was chosen as a balance between excluding direct environmental effects (phenotypic plasticity, environmentally-induced maternal effects) and keeping as much of the original genotypic differentiation as possible (z.e. preventing loss through adaptation to the lab environment). As host plant variation covaried with the latitudinal gradient, our latitudinal patterns could have been confined by patterns of local adaptation to the host plant species. However, we corrected for this potential bias in our analyses, and for all but three traits (diapause incidence, daily fecundity and adult size), no effects of host plant species were found. Only for daily fecundity, where latitude had a significant effect without but not with host plant added to the model, host plant could have masked a pure latitudinal effect. For the other two traits, a latitudinal effect was absent even when excluding host plant from the analysis. In the case of diapause incidence, assessments were made almost immediately after mites were gathered in the field. Therefore, diapause incidence possibly still showed some environmentally induced phenotypic plasticity. Nevertheless, diapause is known to harbor a very strong genetic component (reviewed in Tauber et al. 1986).

By combining an empirical with a detailed, pattern-oriented modeling approach, this study is the first to demonstrate that local adaptation and evolution imposed by the process of range expansion can jointly shape genetic divergence during range expansion along a latitudinal gradient. We were able to show that local adaptation to the growing season mainly affected development time, while the expansion process *per se* induced evolutionary divergence in dispersal and potentially also in intrinsic growth rates. In the current debate on the potential role of local adaptation *versus* phenotypic plasticity during range expansion, our results indicate that local adaptation can effectively drive rapid genotypic changes. It can operate within the same ecological time frame as the process of spatial selection, together thrusting ecological change along the expansion front. To make reliable predictions for expanding populations, we should therefore acknowledge and take into account this fast interplay between both evolutionary forces.

## Online Supplementary material

### S.1: climatological and other information on the field collection sites

**Table S.1.1:**
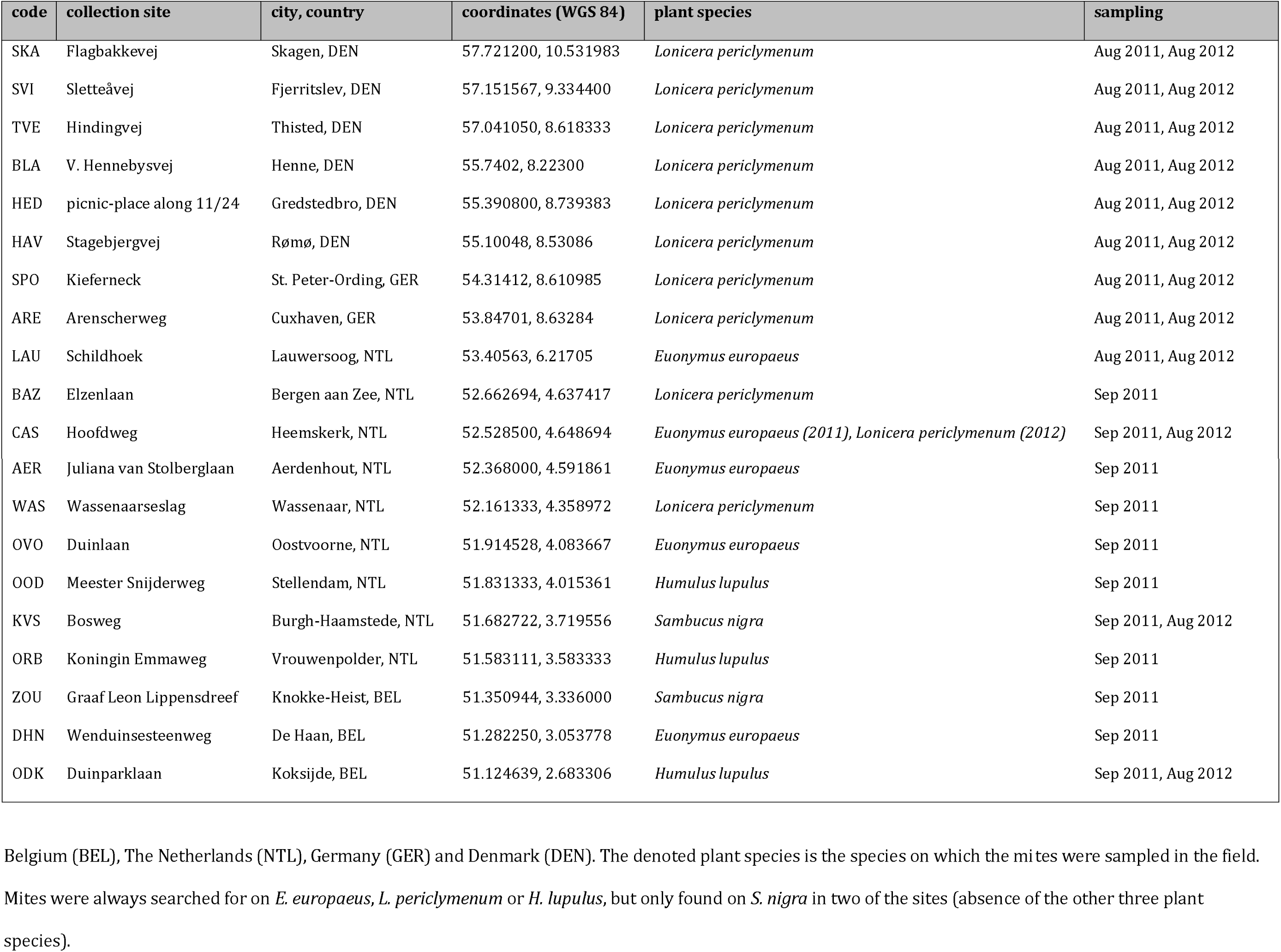
Overview of the field collection sites

**Table S.1.2:** air temperature near surface (land area only) (°C)

|  | January | February | March | April | May | June | July | August | September | October | November | December |
| --- | --- | --- | --- | --- | --- | --- | --- | --- | --- | --- | --- | --- |
| **SKA** | 0,1 | -0,3 | 2 | 5,4 | 10,4 | 14,4 | 15,9 | 15,8 | 12,9 | 9,4 | 5,1 | 2 |
| **SVI** | -0,25064 | -0,3 | 2,04936 | 5,7 | 10,55064 | 14,25064 | 15,7 | 15,6 | 12,7 | 9,14936 | 4,54936 | 1,34936 |
| **TVE** | 0,179 | 0,02537 | 2,12537 | 5,566412 | 10,16204 | 14,021 | 15,30096 | 15,421 | 12,92537 | 9,60437 | 4,984328 | 1,6 |
| **BLA** | 0,801124 | 0,701124 | 2,608062 | 6,3 | 10,80694 | 14,4 | 15,60694 | 15,91388 | 13,50694 | 10,00694 | 5,408062 | 1,915 |
| **HED** | 0,325547 | 0,420112 | 2,620112 | 6,205435 | 10,92011 | 14,32011 | 15,62555 | 15,82555 | 13,21087 | 9,71087 | 5,125547 | 1,725547 |
| **HAV** | 0,5 | 0,6 | 2,9 | 6,4 | 11,2 | 14,5 | 15,9 | 16,1 | 13,4 | 9,9 | 5,3 | 2 |
| **SPO** | 0,6 | 0,9 | 3,361528 | 6,823056 | 11,56153 | 14,86153 | 16,26153 | 16,36153 | 13,6 | 9,9 | 5,3 | 2 |
| **ARE** | 0,7 | 1,170296 | 3,729704 | 7,2 | 11,9 | 15,1297 | 16,6 | 16,6 | 13,8 | 10,0703 | 5,3 | 2,1 |
| **LAU** | 1,61977 | 1,826392 | 4,301309 | 6,986852 | 11,30662 | 14,18685 | 15,9 | 16,1 | 13,6 | 10,01846 | 5,813148 | 3,03954 |
| **BAZ** | 2,552163 | 2,677222 | 4,877222 | 7,477222 | 11,50228 | 14,35012 | 16,27722 | 16,5 | 14,3 | 10,92278 | 6,674941 | 4,027105 |
| **CAS** | 2,547881 | 2,7 | 4,9329 | 7,545802 | 11,58502 | 14,43133 | 16,3329 | 16,5329 | 14,2671 | 10,8671 | 6,627883 | 4,008665 |
| **AER** | 2,617473 | 2,727706 | 5,0584 | 7,634316 | 11,66714 | 14,53581 | 16,43069 | 16,62771 | 14,2 | 10,8 | 6,593389 | 4,112356 |
| **WAS** | 2,921977 | 2,921977 | 5,32697 | 7,563485 | 11,70499 | 14,46349 | 16,56349 | 16,72697 | 14,22198 | 10,82198 | 6,821977 | 4,521977 |
| **OVO** | 3,0998 | 3,101281 | 5,500003 | 7,802567 | 11,90257 | 14,70385 | 16,80257 | 17,00109 | 14,60088 | 11,10088 | 6,998319 | 4,598717 |
| **OOD** | 3,12607 | 3,189992 | 5,5 | 7,927844 | 12,02784 | 14,89177 | 16,92784 | 17,08999 | 14,68999 | 11,21606 | 6,962148 | 4,536078 |
| **KVS** | 3,208275 | 3,308275 | 5,545301 | 8,090603 | 12,0906 | 14,9906 | 16,98233 | 17,2453 | 14,99888 | 11,55358 | 7,108275 | 4,554699 |
| **ORB** | 3,299867 | 3,4 | 5,6 | 8,200133 | 12,20027 | 15,10027 | 17,00027 | 17,30027 | 15,20013 | 11,70013 | 7,2 | 4,499867 |
| **ZOU** | 3,271214 | 3,4 | 5,755787 | 8,228786 | 12,32879 | 15,15757 | 17,03057 | 17,30178 | 15,11721 | 11,61721 | 6,917212 | 4,288426 |
| **DHN** | 3,417733 | 3,4 | 5,8 | 7,982267 | 12,08227 | 14,86453 | 16,66453 | 16,96453 | 14,88227 | 11,28227 | 6,6 | 4,217733 |
| **ODK** | 3,3 | 3,540016 | 5,7 | 8 | 11,81997 | 14,65998 | 16,5 | 16,65998 | 14,6 | 11,2 | 6,680033 | 4,240016 |

**Table S.1.3:**
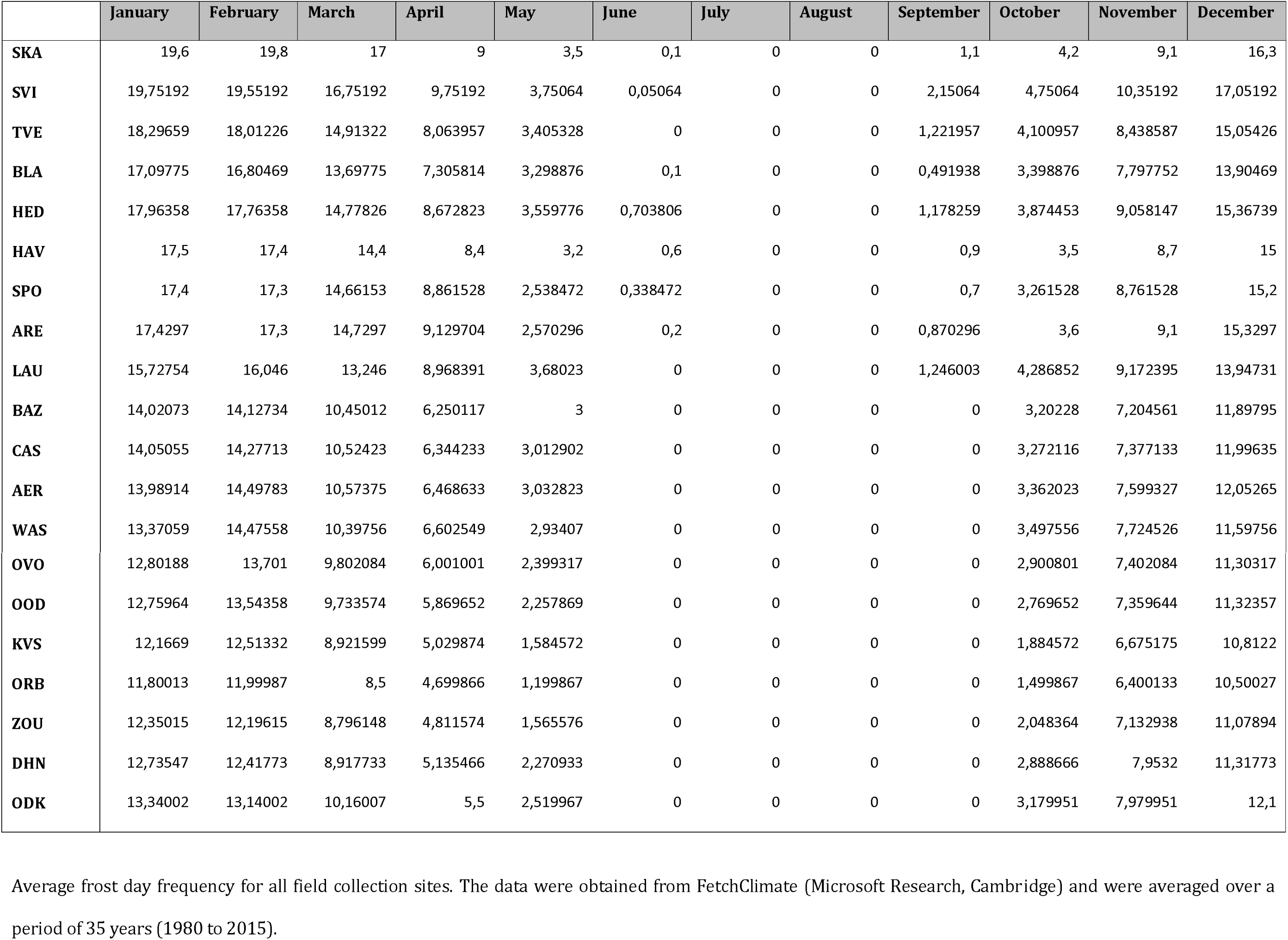
frost days frequency (days/month)

**Table S.1.4:** relative humidity (land only area) (%)

|  | January | February | March | April | May | June | July | August | September | October | November | December |
| --- | --- | --- | --- | --- | --- | --- | --- | --- | --- | --- | --- | --- |
| **SKA** | 91,7 | 89,6 | 87,1 | 82,8 | 81,1 | 81,4 | 82,8 | 83,1 | 84,2 | 87,5 | 89,2 | 90,2 |
| **SVI** | 91,2 | 89,24936 | 86,2 | 80,09872 | 76,99872 | 77,74808 | 79,2 | 80,1 | 83,05064 | 86,95064 | 89,40128 | 90,40128 |
| **TVE** | 90,74881 | 89,60585 | 86,97952 | 81,71215 | 78,68145 | 79,10246 | 79,80341 | 80,30437 | 82,33378 | 85,76544 | 87,80648 | 89,26981 |
| **BLA** | 91,19775 | 89,57919 | 88,09306 | 83,19418 | 79,39531 | 80,20224 | 81,00806 | 81,10112 | 83,10581 | 86,90469 | 88,90357 | 90,80469 |
| **HED** | 92,50218 | 90,27283 | 88,07989 | 82,59076 | 79 | 80 | 81,11467 | 81,1 | 84,58206 | 88,66195 | 90,77663 | 92,40218 |
| **HAV** | 92,4 | 89,9 | 87,7 | 82,3 | 78,9 | 79,9 | 81,1 | 80,9 | 84,3 | 88,2 | 90,4 | 92,5 |
| **SPO** | 93,4 | 89,67694 | 85,95389 | 80,09236 | 77,45389 | 78,45389 | 79,95389 | 79,51542 | 83,77694 | 87,77694 | 90,53847 | 93,46153 |
| **ARE** | 93,25941 | 89,0297 | 84,64059 | 78,71089 | 76,31089 | 77,61089 | 79,04059 | 78,61089 | 83,34059 | 87,2703 | 90,2 | 93,45941 |
| **LAU** | 90,89869 | 88,16167 | 85,79459 | 81,81175 | 79,81175 | 80,28014 | 81,75906 | 81,7406 | 85,36046 | 87,92223 | 89,78685 | 91,31977 |
| **BAZ** | 88,47518 | 86,09111 | 84,3205 | 80,89316 | 79,2886 | 80,04532 | 80,64988 | 80,69772 | 82,31117 | 85,02962 | 86,37974 | 88,52301 |
| **CAS** | 88,60925 | 85,8942 | 84,16208 | 80,61785 | 78,81492 | 79,79863 | 80,56867 | 80,60788 | 82,59662 | 85,28796 | 86,69929 | 88,64847 |
| **AER** | 88,60236 | 85,49642 | 83,90878 | 80,21095 | 78,28749 | 79,55762 | 80,35187 | 80,38768 | 82,87674 | 85,49209 | 86,90531 | 88,66289 |
| **WAS** | 87,82452 | 84,92697 | 83,74895 | 79,88546 | 78,46094 | 79,70744 | 79,61698 | 79,86349 | 82,58801 | 84,99301 | 86,09301 | 88,11498 |
| **OVO** | 87,49932 | 84,99763 | 83,69526 | 79,68973 | 78,38933 | 79,38993 | 79,49466 | 79,59378 | 82,19694 | 84,59863 | 85,89972 | 88,001 |
| **OOD** | 87,38394 | 84,87216 | 83,44431 | 79,26683 | 77,9929 | 78,94076 | 79,29645 | 79,28039 | 81,95609 | 84,36788 | 85,75787 | 87,92179 |
| **KVS** | 87,11767 | 84,92595 | 83,28065 | 79,32426 | 78,27784 | 78,81599 | 79,45189 | 78,88177 | 81,4641 | 83,64755 | 85,24642 | 87,40112 |
| **ORB** | 87,09987 | 84,99973 | 83,29947 | 79,3992 | 78,49947 | 78,79934 | 79,59946 | 78,69933 | 81,2996 | 83,29974 | 85,1 | 87,19986 |
| **ZOU** | 87 | 84,81543 | 83,14064 | 79,10028 | 78,84865 | 79,35843 | 80,03322 | 78,77922 | 81,04421 | 83,12878 | 85,12878 | 87,04421 |
| **DHN** | 87,01773 | 84,91773 | 83,6532 | 79,6532 | 80,22413 | 81,04187 | 81,3064 | 79,58867 | 81,01773 | 83,1 | 85,1 | 87,1 |
| **ODK** | 87,90008 | 85,08003 | 83,11997 | 78,41997 | 80,11997 | 81,8 | 81,75998 | 81,3401 | 81,72005 | 83,86006 | 86,72013 | 88,62013 |

**Table S.1.5:** sunshine fraction (% of maximum possible sunshine)

|  | January | February | March | April | May | June | July | August | September | October | November | December |
| --- | --- | --- | --- | --- | --- | --- | --- | --- | --- | --- | --- | --- |
| **SKA** | 12,4 | 20,7 | 29,4 | 39,5 | 44,5 | 46,1 | 46 | 45,6 | 37,6 | 28,3 | 21,4 | 13,9 |
| **SVI** | 14,29872 | 23,6 | 30,5 | 41,44936 | 46,44936 | 46,2 | 46,44936 | 46,4 | 38,25064 | 29,25064 | 23,2 | 15,5 |
| **TVE** | 14,99615 | 24,04785 | 30,69178 | 41,89178 | 46,52344 | 46,19415 | 46,18785 | 46,24733 | 37,4147 | 28,6727 | 22,6937 | 15,39274 |
| **BLA** | 16,05818 | 24,55124 | 30,85593 | 39,54205 | 44,63512 | 41,36644 | 39,94675 | 43,55368 | 34,96531 | 27,88612 | 21,07337 | 14,98725 |
| **HED** | 14,92392 | 23,21848 | 29,28532 | 38,46902 | 44,37663 | 41,36358 | 40,04728 | 43,86195 | 35,07826 | 28,39837 | 20,87445 | 15,09837 |
| **HAV** | 15,4 | 23,6 | 29,2 | 38,2 | 44,2 | 41 | 39,5 | 43,6 | 34,7 | 28,4 | 20,5 | 15,1 |
| **SPO** | 16,56153 | 24,76153 | 28,83847 | 38,43847 | 44,57694 | 41,5 | 39,73847 | 44,5 | 35,36153 | 29,26153 | 20,23847 | 15,7 |
| **ARE** | 16,65941 | 25,0297 | 28,5 | 37,95941 | 43,68911 | 40,84852 | 39,21881 | 44,11882 | 35,18911 | 29,4297 | 20,2297 | 15,2297 |
| **LAU** | 15,81315 | 24,63423 | 29,04607 | 37,14607 | 41,03944 | 39,25791 | 36,44607 | 40,62629 | 34,33823 | 28,5277 | 20,19338 | 14,41846 |
| **BAZ** | 17,89772 | 26,59772 | 31,341 | 39,17266 | 44,57038 | 42,09749 | 40,25649 | 43,32482 | 36,29544 | 30,37722 | 20,64556 | 16,2 |
| **CAS** | 17,79498 | 26,54788 | 31,04416 | 39,05577 | 44,35075 | 41,83942 | 40,26945 | 43,22365 | 36,12996 | 30,4329 | 20,58 | 16,2 |
| **AER** | 17,63648 | 26,61747 | 30,58025 | 38,90067 | 44,06636 | 41,49623 | 40,34884 | 43,18532 | 35,99193 | 30,47229 | 20,50512 | 16,14309 |
| **WAS** | 17,39001 | 26,99046 | 29,92608 | 38,64107 | 43,84606 | 41,03563 | 40,36804 | 43,18002 | 35,96549 | 30,2485 | 20,59789 | 15,65805 |
| **OVO** | 17,50068 | 27,20149 | 29,7986 | 38,29832 | 43,29763 | 40,59732 | 40,1996 | 43,30029 | 36,70522 | 30,60433 | 21,60494 | 15,80157 |
| **OOD** | 17,61606 | 27,18571 | 29,90428 | 38,26215 | 43,14609 | 40,66643 | 40,22607 | 43,38999 | 37,07175 | 30,90783 | 21,82962 | 16,03212 |
| **KVS** | 17,99888 | 26,79172 | 30,52482 | 38,36297 | 42,64755 | 41,30827 | 40,45358 | 43,67952 | 37,6359 | 31,50715 | 22,00827 | 16,80715 |
| **ORB** | 18,20013 | 26,7 | 30,8 | 38,4 | 42,29973 | 41,40013 | 40,60027 | 43,89987 | 37,8004 | 31,8004 | 22,10013 | 17,10027 |
| **ZOU** | 18,51157 | 27,68515 | 32,13916 | 38,99258 | 42,29822 | 41,83679 | 41,24836 | 45,72816 | 38,45015 | 33,06973 | 23,98752 | 17,82136 |
| **DHN** | 18,7 | 29,12413 | 34,17733 | 39,77093 | 42,7532 | 42,4532 | 42,07093 | 48,90146 | 38,81773 | 34,3064 | 26,63053 | 18,3532 |
| **ODK** | 19,12005 | 29,77995 | 35,01997 | 40,3 | 43,18003 | 43,08003 | 42,7 | 50,19992 | 39,72005 | 35,34002 | 27,91997 | 18,94002 |

Average sunshine fraction for all field collection sites. The data were obtained from FetchClimate [Microsoft Research, Cambridge) and were averaged over a period of 35 years [1980 to 2015).

### S.2: Used methodology for data collection and statistics

#### Dispersal propensity and latency

Dispersal propensity and dispersal latency of mites from 2012 were assessed using a similar setup as described in Van Petegem et al. (2015). During a three hour observation in a climate room at 27°C, aerial dispersal propensity was assessed by counting the percentage of female mites showing the aerial take-off posture (*i.e*. upraised first pair of legs and cephalothorax, to increase drag). Dispersal latency was then assessed by counting the number of minutes between the start of the three hours of observation and the moment the focal female showed the take-off posture. Only one-day-old, freshly mated females were used, since they are the main dispersing stage (see Li and Margolies 1994; Van Petegem, Petillon et al. 2015 for more details). The experiment was repeated for different actual and maternal densities since dispersal is known to be a density-dependent process in which genotypes might have different density sensitivities and thresholds (Bitume, Bonte et al. 2014). High, medium and low densities were obtained by performing the synchronization (see earlier) on a piece of leaf of, respectively, 4cm^2^, 7.5cm^2^ or 12cm^2^. This gave a measure of the density experienced by the mother at the time of egg laying (*i.e*. maternal density). The density experienced by the focal mites themselves was also measured by quantifying the density of female offspring on each piece of leaf (*i.e*. actual density). Because of the obvious strong correlation between both density measures, only the most explanatory measure (for the variation in the dispersal data) was later kept for the statistical analysis.

#### Diapause incidence

Between nineteen and twenty-five (Danish and German populations) and six and eleven (Dutch and Belgian populations) days after sampling a population in the field (in 2011), data on diapause incidence of the population were collected. At this point, the mites were still temporarily stored on bean leaves in a regular room with no specific light regime. Mites had spontaneously started to go into diapause, allowing us to assess the proportion of diapausing female mites for each population.

#### Fecundity and longevity

Fecundity and longevity of mites from 2011 were assessed by putting a one-day-adult female on a small piece of bean leaf and counting the number of days and the number of eggs per day until she died. Data on both daily and lifetime fecundity were thus obtained. For each population, three same-sized pieces of bean leaf were put on wet cotton in each of four Petri dishes, resulting in twelve replicates per population. All Petri dishes were stored in a climate room at 27°C, with a light-regime of 16:8 LD.

#### Egg and juvenile survival and development time

Egg survival, juvenile survival and development time of the mites from 2011 were assessed by putting three four-day-adult female mites on a piece of bean leaf and following up the development of their offspring. Four-day-adult females were used because, on average, female *T. urticae* have their maximum daily fecundity when they are four to five days old (own observation). The females were allowed to lay eggs for 24 hours in a climate room at 20°C, with a light-regime of 16:8 LD, after which they were removed and their eggs were left to develop. We here used a temperature of 20°C because this allowed a steady development, hence a higher probability of detecting differences in development time. (The temperature of 27°C, used in all other setups, was chosen to allow a relatively fast development and therefore a relatively fast progress of the experiments). For each population, three same-sized pieces of bean leaf were put on wet cotton in each of four Petri dishes, resulting in twelve replicates per population. All Petri dishes were checked daily at approximately the same hour of the day to examine (i) for each egg if it had hatched (ii) the developmental stage of each juvenile (iii) the sex of each freshly molted adult. Afterwards, egg survival was calculated as the number of hatched eggs divided by the total number of eggs, juvenile survival as the number of adults divided by the number of larvae, and development time as the number of days between the day an egg was laid and the molt into an adult spider mite (*i.e*. age at maturity).

#### Sex ratio

Offspring sex ratio was assessed in all populations from 2012 by putting a one-day-adult, freshly mated female on a small piece of bean leaf and allowing her to lay eggs during seven days, after which the sex of her offspring could be determined. (We chose seven days because the majority of eggs is laid within this period (Krainacker and Carey 1989).) More specifically, sex ratio was calculated as the number of male offspring divided by the total number of offspring. For each population, three same-sized pieces of bean leaf were put on wet cotton in each of five Petri dishes, resulting in fifteen replicates per population. All Petri dishes were stored in a climate room at 27°C, with a light-regime of 16:8 LD.

#### Adult size

For each population of 2012, between twenty-seven and thirty two-day-adult females were immobilized through snap freezing at -80°C and later photographed one by one with a digital camera (Nikon Coolpix 4500) mounted on a stereomicroscope. To be able to calibrate the photographs, each female was positioned on a small measuring plate (accurate to 50μm). Using ImageJ 1.47v (Wayne Rasband, National Institutes of Health, USA), all photographs were afterwards analyzed and the surface area (mm^2^) of the mites (legs and capitulum excluded) was calculated (Schneider, Rasband et al. 2012).

#### Statistics

Prior to univariate analyses, we performed a multivariate distance-based ANOVA to test for variation in multivariate life history parameter space using the vegan and permute packages of R version 3.1.0 (The R Foundation for Statistical Computing, 2014). Relationships between Gower’s distances of all life-history traits were assessed with the Adonis function. Because our trait quantification induced different error structures for the different traits, we used overall population averages. As this ANOVA showed considerable variation in life-history strategies among the different sampled populations (F_1,8_=2.2285; p=0.051 for the subset of 12 populations sampled in both 2011 and 2012, and hence all nine measured traits; and F_1,16_=3.6568; p=0.009 for eighteen of the twenty populations sampled in 2011 and hence the subset of six traits measured for only 2011), we then performed univariate analyses.

All univariate analyses were performed in SAS 9.4 (SAS Institute Inc. 2013), using the MIXED procedure for linear mixed models (analysis of dispersal latency, lifetime and daily fecundity, longevity, development time and adult size) or the GLIMMIX procedure for generalized linear mixed models (analysis of dispersal propensity, diapause incidence, egg and juvenile survival and sex ratio). Latitude and host plant species (*i.e*. the plants species from which mites were gathered in the field) were always used as the independent variables, and for the analysis of dispersal propensity and latency, maternal density and its interaction with latitude were also added. Finally, for the analysis of diapause incidence, the time lag between the collection of mites in the field and the observation of the behavior in the lab was added as a covariate. For all generalized linear mixed models, a binomial error structure was modeled with the proper link function. Petri dish identity (nested within population) was furthermore modeled as a random effect to control for dependency among the leaves on each Petri dish (except for dispersal and adult size, where no Petri dishes were used). For egg and juvenile survival and for development time, leaf identity-nested within Petri dish-was additionally modeled as a second random factor to control for dependency among the mites on each piece of leaf. By modeling residual variation as an additional random factor (in the glimmix procedure), we corrected for potential overdispersion (Verbeke and Molenberghs 2000). The denominator degrees of freedom for the tests of fixed effects were computed according to a general Satterthwaite approximation. All non-significant contributions (p>0.05) were omitted by a backwards selection procedure. Finally, post-hoc Tukey tests were used to obtain the pair-wise differences among densities (analysis of dispersal propensity and latency) and host plant species.

Additionally, we tested whether the data on development time followed a saw-tooth pattern by fitting a fourth order polynomial to these data. This was done in SAS 9.4 (SAS Institute Inc. 2013), using the MIXED procedure. In order to obtain a simple, but solid model, population averages of development time were used. Latitude was furthermore used as the independent variable and gender as a covariate (to account for differences in development time between male and female mites). The denominator degrees of freedom for the tests of fixed effects were computed according to a general Satterthwaite approximation.

### S.3: Settings of the individual-based model

#### S.3.0: main text

##### spatial dimensions and dispersal

The model landscape consisted of one hundred rows (latitude) and five columns (longitude) of 10km by 10km patches (grid cells). All patches were occupied from the start in the stable range scenario, while only the ten southernmost patches were initially occupied in the two range expansion scenarios. Mites were allowed to disperse from one patch to a neighbouring patch, but with a 90% dispersal mortality. Only adult individuals could disperse and their probability to do so was embedded in their genome as an emigration rate. This rate underwent a mutation probability of 1% at the birth of an individual, with an effectsize sampled from a uniform distribution between -10% and +10%. We did not include density-dependent dispersal, as this would not make sense given the used scale (grid cells much bigger than the size of a single leaf or plant-where density dependence would occur) and the low population sizes in our model (see further on).

##### temporal and spatial temperature gradients

To simulate appropriate local temperatures throughout the year, two trigonometric functions where created to match the actual average daily temperatures in locations in the extreme south and north of our study area. (The used weather stations were, respectively, Brussels, Belgium and Gothenburg, Sweden). For locations in between these two extremes, an intermediate function was calculated, assuming that the two functions converged linearly. This resulted in a unique temperature regime for each latitude, and an accompanying gradient in the length of the growing season.

##### population dynamics

Using data from Sabelis (1981), the development time (see supplementary material S.3.1), longevity (see supplementary material S.3.2), juvenile and adult mortality rate (see supplementary material S.3.3), and lifetime fecundity (see supplementary material S.3.4) of the mites in the model were simulated according to their locally experienced temperature. The mean daily fecundity for mites at a certain temperature could then be calculated by dividing their expected lifetime fecundity by their expected longevity at this temperature (see supplementary material S.3.4). The actual simulated number of produced eggs for a specific individual mite could subsequently be sampled from a normal distribution with this calculated mean and a standard deviation of 1.48, which was based on empirical data from another study on *T. urticae* (A. De Roissart, unpublished data). Since we only consider females in our model, we assumed asexual reproduction and therefore took only 75% (cfr. sex ratio 3:1) of the number off eggs produced, to take into account the absence of males (Sabelis 1981).

Each time step, all individuals faced a temperature-dependent mortality which increased rapidly as local temperatures reached either too low (10°C) or too high (35°C) values (see supplementary material S.3.3) and which was absolute when temperatures dropped below 10°C. To avoid this temperature-dependent mortality, adult spider mites in the model could go into diapause. They then, however, aged slower and could not produce eggs. The diapause behavior of the mites was determined by two loci subject to selection/mutation. One locus determined the temperature for which mites entered diapause in ‘autumn’ and the other locus determined the temperature for which mites terminated diapause in ‘spring’. To nonetheless account for extreme weather conditions in winter, we each winter imposed a mortality event, whereby 50% of all diapausing individuals in the south and 80% of all diapausing individuals in the north were killed (with a linear increase in winter mortality inbetween the extreme south and north).

At the end of each time step, local population sizes were assessed for all patches. If overcrowding occurred, individuals were randomly deleted from the patch until the number of individuals matched the carrying capacity (K=200). This procedure improved computational performance by keeping populations at a manageable size. As a consequence, however, the population sizes in our model were relatively low given the large spatial scale. We compensated for this by assuming a relatively low dispersal mortality (90%, see earlier). This compromise was unavoidable, as it is impossible to combine realistic density-dependence (on the level of a single plant or leaf) with the large spatial scale required to investigate range expansion dynamics.

##### trade-offs

Several trade-offs were implemented in the model, by attributing three specific traits to the mites. These traits were embedded in the genome as a single locus, subject to a mutation rate of 1% at the birth of an individual. A mutation’s effectsize was sampled from a uniform distribution between -10% and +10%.

The first two traits concerned diapause behavior: the temperatures at which mites decided to enter diapause in autumn and to reactivate in spring. Postponing diapause in autumn allowed continued egglaying, but came with the cost of an increased temperature-dependent mortality risk. Terminating diapause sooner (*i.e*. at lower temperatures) in spring came with the competitive benefit of producing eggs more early in the season, but with the same mortality risk.

As a third trait, we allowed mites to invest in a faster development at the expense of their fecundity and *vice versa*. Such a trade-off has to date not been found in *T. urticae*, but is likely to exist. The development time in this mite species is just so short, that small changes in it would easily go unnoticed, unless mites are monitored constantly. In fact, not a single true life-history trade-off has so far been found in *T. urticae* (*e.g*. Li and Margolies 1994; Bitume, Bonte et al. 2011; Tien, Sabelis et al. 2011; Fronhofer, Stelz et al. 2014). Yet, trade-offs are expected to exist in this species and were truly essential in our model, as mites would otherwise evolve to become ‘Darwinian demons’ which can maximize all aspects of their fitness simultaneously. We thus choose a trade-off which has been observed in several species: a trade-off between development and fecundity (*i.e*. a longer development time, hence slower development, results in an increased fecundity) (e.g. Nunney 1996; Tsikliras, Antonopoulou et al. 2007; Lewis, Brakefield et al. 2010; Yadav and Sharma 2014). This trade-off can be mediated through a positive correlation between development time and adult size (*e.g*. Nunney 1996) and was coded as a single value varying between 0 and 1 (see supplementary material S.3.5). This single locus trait could evolve under selection and resulted in a deviation (increase or decrease) in the investment in fecundity or development from the one expected based on the local temperature (see supplementary material S.3.1 and S.3.4). These deviations were limited to 10 or 20% of the values expected under this current local temperature. A value of the trade-off locus of 0 meant maximal investment in development (*i.e*. in a fast development, hence a short development time) at a maximal cost of fecundity, while a value of 1 imposed the opposite. Logically, a value of 0.5 thus did not result in any shift in investment. Importantly, a ‘maximal’ shift in development was not necessarily of the same magnitude as a ‘maximal’ shift in fecundity (see supplementary material S.3.5 for a detailed explanation). When comparing our empirical results with the simulation results, we therefore conducted an additional analysis to see what trade-off settings (*i.e*. which of all possible combinations of 10 and 20%, see supplementary material S.3.5) resulted in the best fit with our empirical data and were thus most realistic.

#### S.3.1: Development as a function of temperature

The development of juvenile spider mites depends on the local temperature and lasts about nine days under optimal conditions (±28°C, see Sabelis 1981). We assessed the effect of temperature on the daily progress in development (E), using data by Sabelis (1981). Our calculations were based on formulas which were developed by Logan (1976) and Lactin (1995), and which were used by Bancroft & Margolies (1999) to determine how well development is progressing under certain temperatures (*e.g*. E=1 is an optimal development; E=0.5 is 50% slower). The fit between the data from Sabelis (1981) and our predicted values was high (R^2^ = 0.9893). E was calculated as:

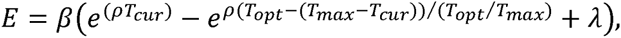
where *β* [0–1) is used to scale the development rate; *ρ* is the constant growth multiplier below the optimal temperature; T_max_ is the maximum temperature at which development can take place [38.1°C); T_cur_ is the local ambient temperature; T_opt_ is the optimal temperature for development [28.17°C); and λ [-1.74) allows the curve to intersect the abscissa atsuboptimal temperaturs [Bancroft and Margolies 1999).

Each day, the daily progress in juvenile development (E) was calculated and added to the value so far (z.e. the sum of all E-values of the previous days). When this summed E-value surpassed 7 (*e.g*. after seven days under optimal conditions), the individual became an adult.

**Figure S.3.1:**
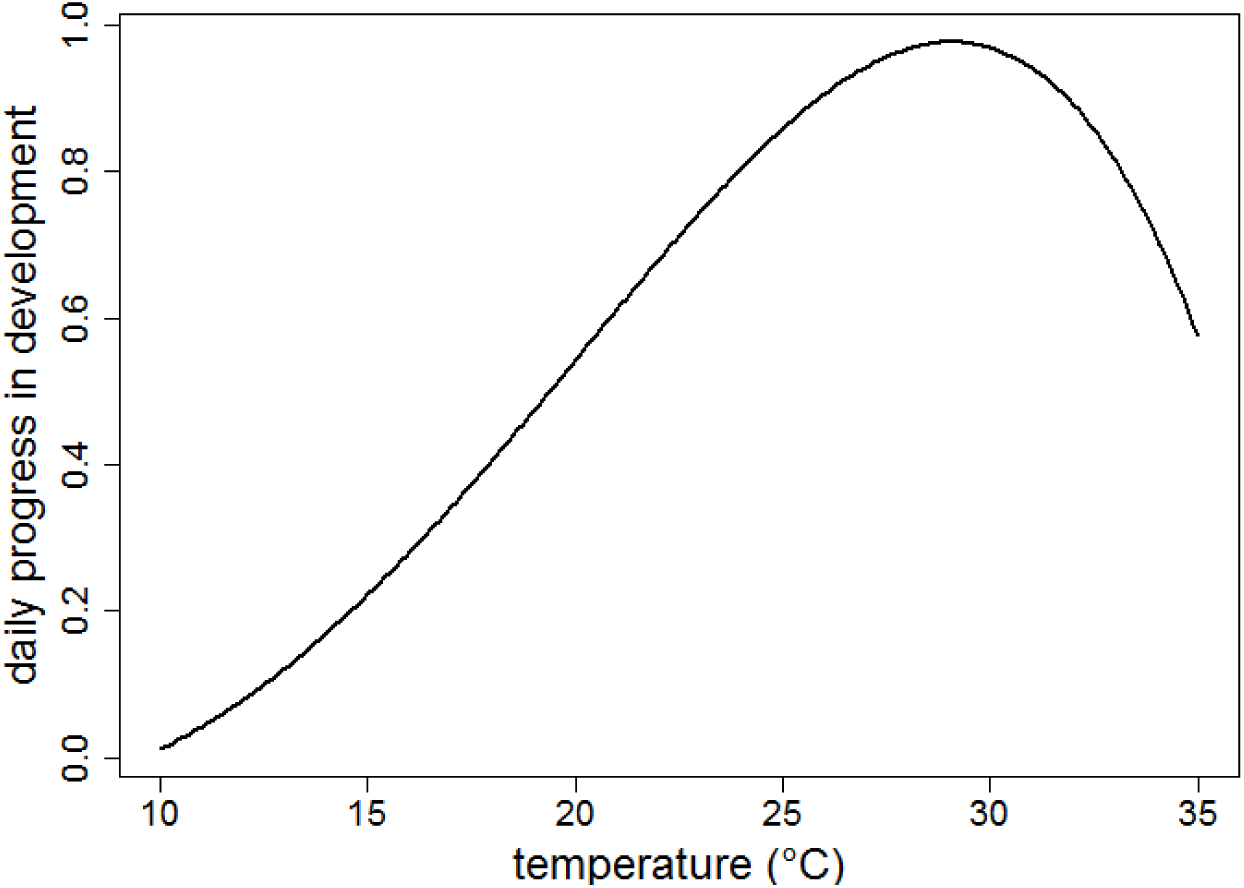
predicted development as a function of temperature

#### S.3.2: Longevity as a function of temperature

Using the data from Sabelis (1981), we predicted the lifetime expectancy for an adult mite (N_days_), depending on the local temperature. The fit between the data from Sabelis (1981) and our predicted values was high (R^2^ = 0.9779):

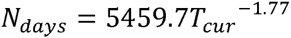

**Figure S.3.2:**
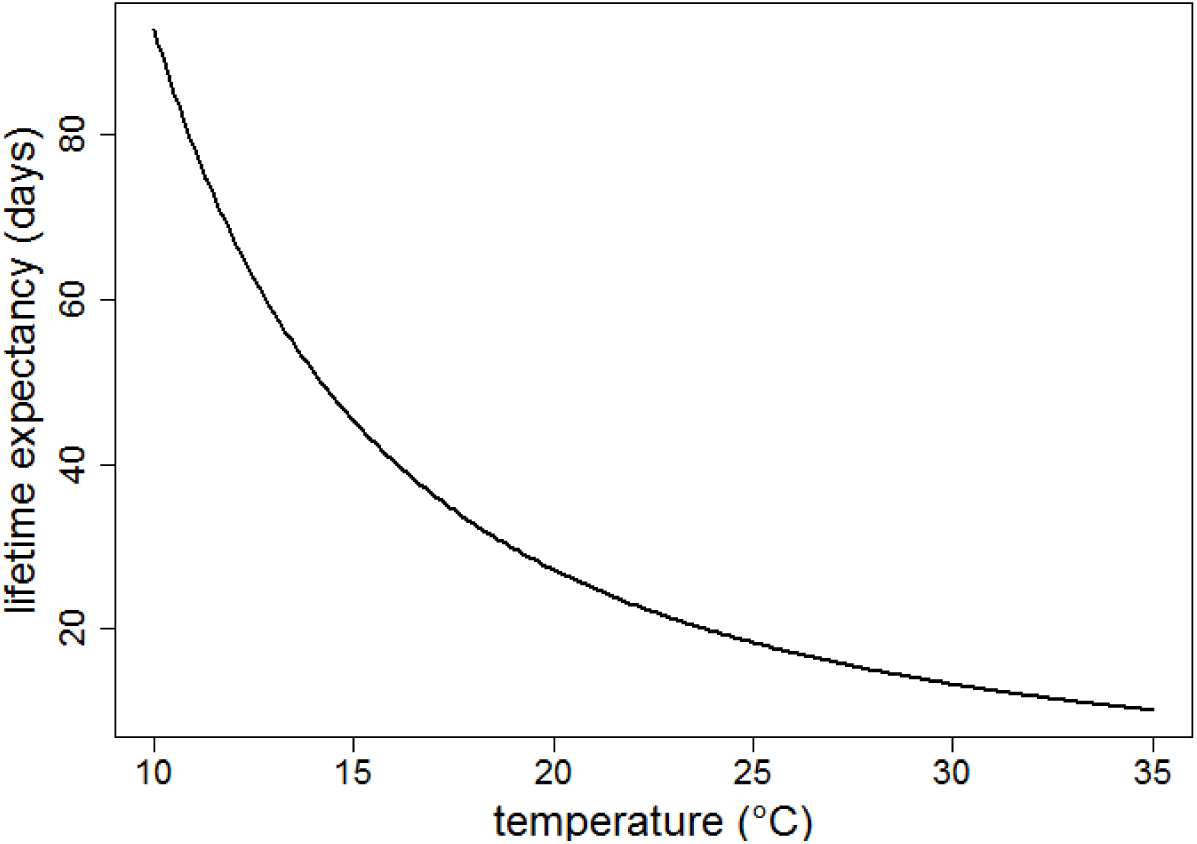
predicted longevity as a function of temperature

#### S.3.3: Juvenile and adult mortality as a function of temperature.

To assess the mortality caused by suboptimal temperatures (a mortality additional to the mortality due to age, which also depends on temperature -see S.4.2), we fitted nonlinear functions to the data from Sabelis (1981). We did this for both juveniles and adults:juvenile temperature-dependent mortality function (fit between used data from Sabelis (1981) and our predicted values: R^2^ = 0.9996):

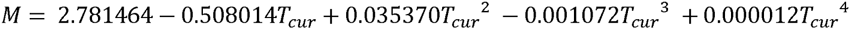
adult temperature-dependent mortality function (fit between used data from Sabelis (1981) and our predicted values: R^2^ = 0.9996):

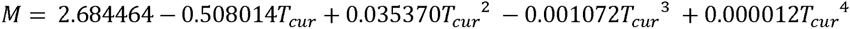

To assess the daily probability of a temperature-induced death, we divided the mortalities for each stage (juvenile or adult) by the presumed length of that stage given the current local temperature.

**Figure S.3.3:**
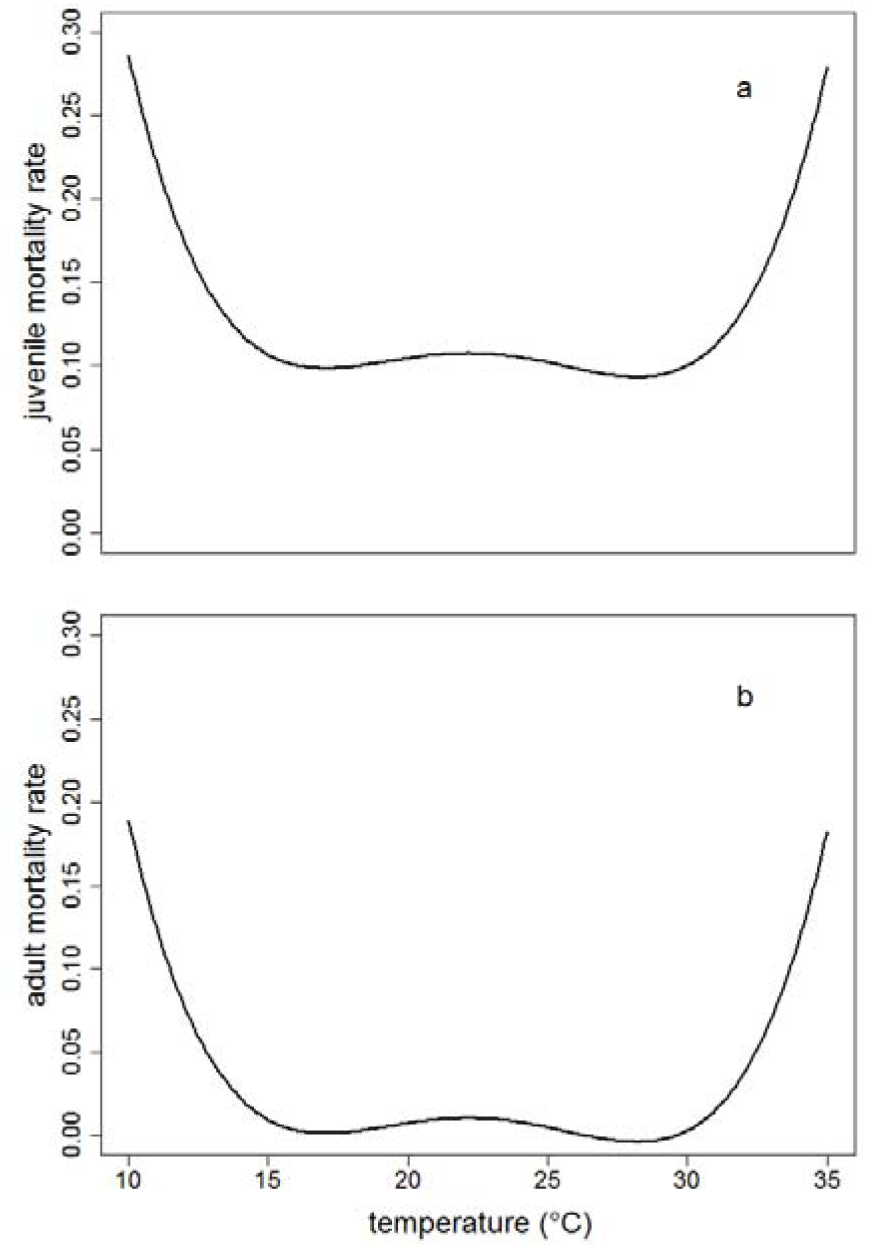
predicted juvenile (a) and adult (b) mortality rate as a function of temperature

#### S.3.4: Fecundity as a function of temperature

We estimated the (potential) number of eggs produced during a mite's lifetime (N_e_), depending on the current local temperature. We therefore fitted nonlinear functions to the data from Sabelis (1981).The fit between these data and our predicted values was high ( R^2^ = 0.9777).

**Figure S.3.4:**
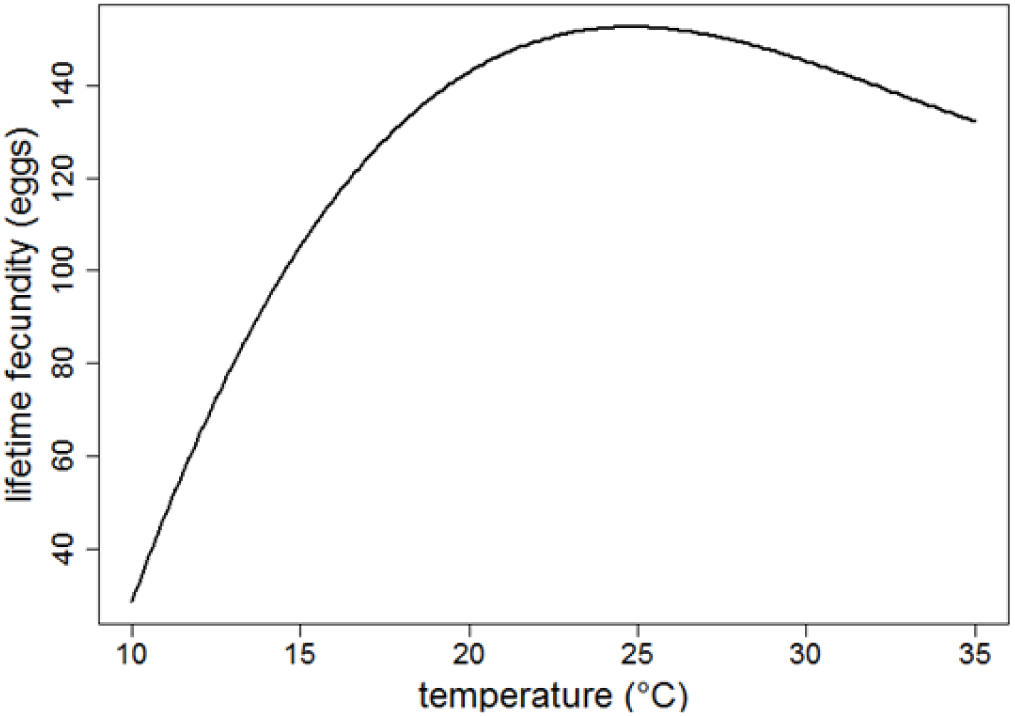
predicted lifetime fecundity as a function of temperature

To assess the daily fecundity, we divided the estimated lifetime fecundity by the predicted longevity of the mite (with both these traits depending on the local temperature).

#### S.3.5: Trade-off balance between development and fecundity

**Figure S.3.5:**
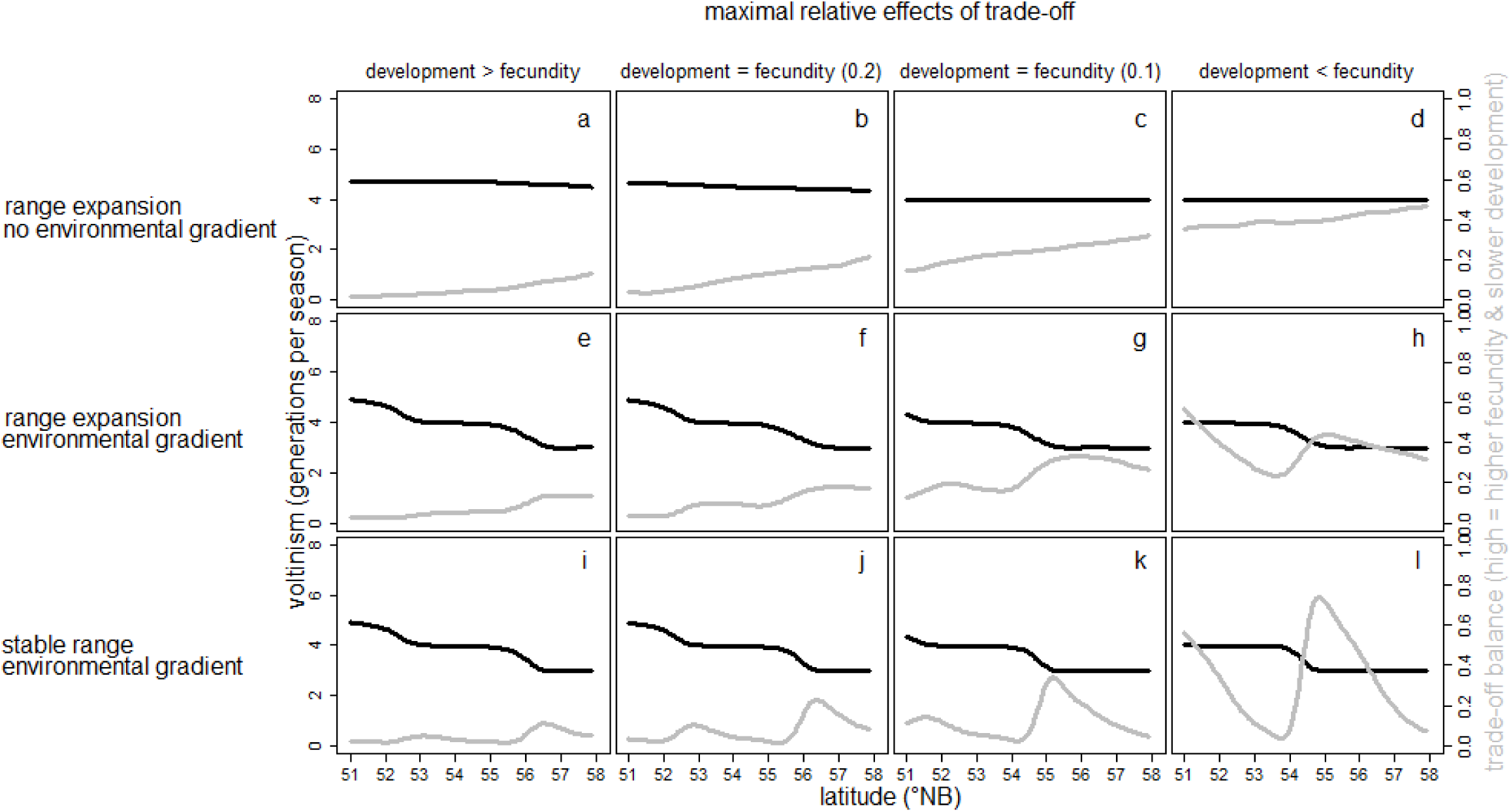
An overview of how the maximal trade-off effect on development *vs*. fecundity affects voltinism (black line) and the trade-off balance between development and fecundity (grey line) along the latitudinal gradient.

The four trade-off settings vary the maximal costs and benefits for development vs. fecundity. In the first column, the trade-off effect on development (*max*. ±20%) is larger than on fecundity (*max*. ±10%). In the second column, both trade-off effects are equal and relatively large (*max*. ±20%). In the third column, both trade-off effects are equal and relatively small (*max*. ±10%). In the fourth column, the trade-off effect on development (*max*. ±10%) is smaller than on fecundity (*max*. ±20%). The trade-off function between both traits is linear. In other words, when the trade-off balance [grey line) is 0.5, there is no effect on either trait When the trade-off balance is 1, however, there is a maximal investment in fecundity at a maximal expense of development [e.g. +10% for fecundity and +20% for development time, thus a 20% slower development, in case of the first column). For a trade-off balance of 0, the opposite is true: a maximal investment in development at a maximal expense of fecundity [e.g. -20% for development time, thus a 20% faster development, and -10% for fecundity in case of the first column). The effects are shown for our three model scenarios [one scenario with range expansion in a homogeneous landscape, one scenario with range expansion along an environmental gradient and one scenario with a stable range along this environmental gradient).

### S.4: Statistical patterns used to compare model and empirical data

We compared our empirical and model data through statistical patterns of those three traits that were subject to spatial and natural selection and for which a comparison between simulation and empirical data could be made: the regression slope of intrinsic growth rate, dispersal, and develoment time, and the amplitude and wavelength of the teeth in the saw-tooth pattern in development time. The slopes were each time calculated from a linear regression over the latitudinal gradient The comparison of the empirical and simulated saw-tooth pattern was less straightforward. Under certain model settings, a clear latitudinal saw-tooth pattern in the trade-off balance between development time and fecundity emerged (see figure S.3.5). From this simulated latitudinal pattern in the trade-off balance, we could calculate a latitudinal pattern for development time. To be able to compare this calculated pattern in development time with the saw-tooth pattern found for our empirical data, we then assessed local development times along the simulated latitudinal gradient, as if an optimal temperature was present at all latitudes (*cfr*. the empirical assessment of development time: performed in common garden at optimal rearing conditions). More particular, a comparison between the calculated and the empirical sawtooth pattern was made through a comparison of two parameters: the amplitude and the wavelength of the saw-theeth. These parameters could be assessed by fitting a smoothing spline to both the calculated and the empirical saw-tooth pattern. The average amplitude was then each time assessed by taking the mean difference between subsequent maxima and minima in development time.The average wavelength, on the other hand, was assessed by each time taking the mean distance (in latitude) between subsequent maxima and subsequent minima in development time.

### S.5: Life-history trait variation along the latitudinal gradient

Latitude did not affect diapause incidence (figure S.5a), juvenile survival (figure S.5b) or adult size (figure S.5c). Diapause incidence and adult size, together with daily fecundity, were instead affected by the host plant species on which mites were collected in the field. Mites originating from *L. periclymenum* (33.15 ± 9.50 SE) had a (marginally) significantly higher diapause incidence than those originating from *H. lupulus* (0.19 ± 0.18 SE) (t_46,57_=-4.75; p<0.0001), *S*. *nigra* (0.24 ± 0.26 SE) (t_75,32_ = 4.20; p=0.0006) or *E. europaeus* (5.23 ± 2.93 SE) (t_25,04_=2.59; p=0.0585). The difference in diapause incidence between mites collected on *E. europaeus* and *H. lupulus* (t_64,35_=-3.36; p=0.0079) or *S. nigra* (t_126.1_=2.83; p=0.0331) was also significant Mites originating from *L. periclymenum* (0.078 ± 0.001SE) were moreover significantly larger than those from 5. *nigra* (0.072 ± 0.002SE) (t_343_= 3.00; p=0.0153). Furthermore, mites collected on *L. periclymenum* (3.82 ± 0.32SE) laid significantly less eggs than mites collected on *H. lupulus* (5.69 ± 0.49SE) (t_69_=3.18; p=0.0115), *S. nigra* (5.80 ± 0.60SE) (t_69_=-2.91; p=0.0248) or *E. europaeus* (5.19 ± 0.35SE) (t_69_=-2.90; p=0.0249).

**Figure S.5:**
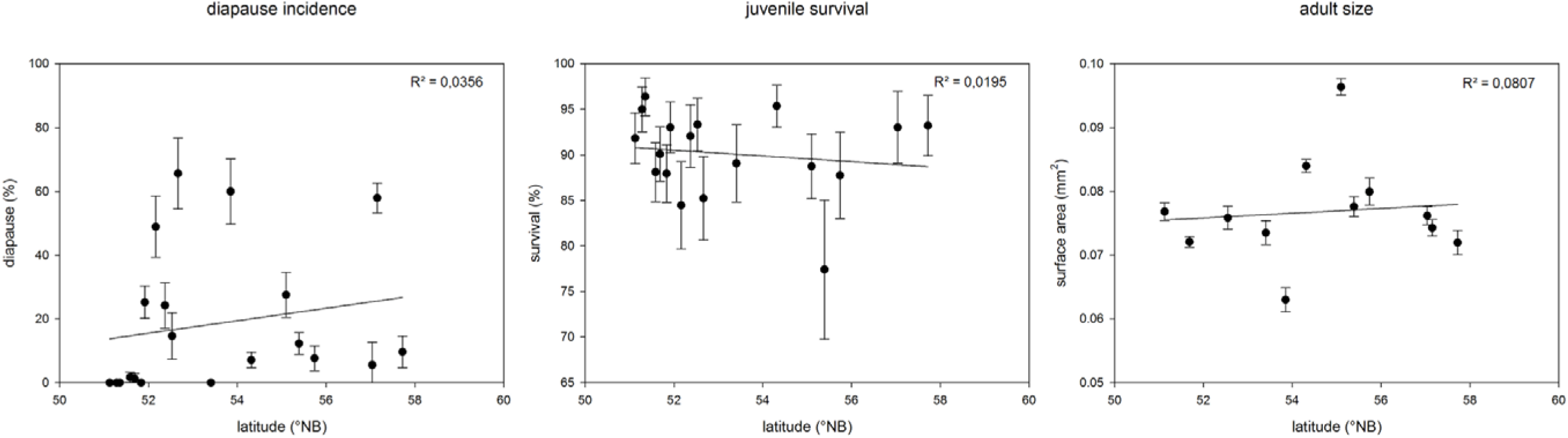
diapause incidence, juvenile survival and adult size as a function of latitude. Diapause incidence (a), juvenile survival (b) and adult size (c) for each sampled population along the latitudinal gradient. Population means are given ± 1 standard error (bars). Regression lines are shown together with there R^2-^values. These values are calculated in a conservative way, using the population averages.

### S.6: Patterns emerging from the model

Due to a number of assumptions (*e.g*. the strength of the trade-off between development and fecundity), we had to simulate a large parameter space. Nonetheless, three consistent (*i.e*. consistent over all the trade-off settings) patterns emerged. The first emerging pattern was the presence of a steap increase in dispersiveness from south to north in both the expansion scenarios, while this increase was absent under the stable range scenario (figure S.6a). The second pattern was a clear stepwise decrease in the number of generations per season (voltinism) from south to north in the scenarios with an environmental gradient (figure S.6b, grey line). The third pattern concerned the temperatures at which mites terminated diapause in spring and entered diapause in autumn. In all model scenarios, the former temperature consistently evolved to lower values than the latter (figure S.6b, black lines). Most probably, this is because eggs laid at the very end of the season will not have the time to mature, while the very first eggs of the season are important for population growth. Therefore, entering diapause later in autumn can be expected to be less advantageous than terminating diapause earlier in spring, resulting in a higher evolutionary pressure on the latter. Under conditions where a step-wise decrease in voltinism occurred from southern to northern latitudes, a pronounced saw-tooth pattern moreover emerged in the temperature at which mites terminated diapause (figure S.6b, dotted black line) but not in the temperature at which they entered diapause. (Note that this saw-tooth pattern was not a consistent pattern, as it did not occur under all trade-off settings.)

**Figure S.6a:**
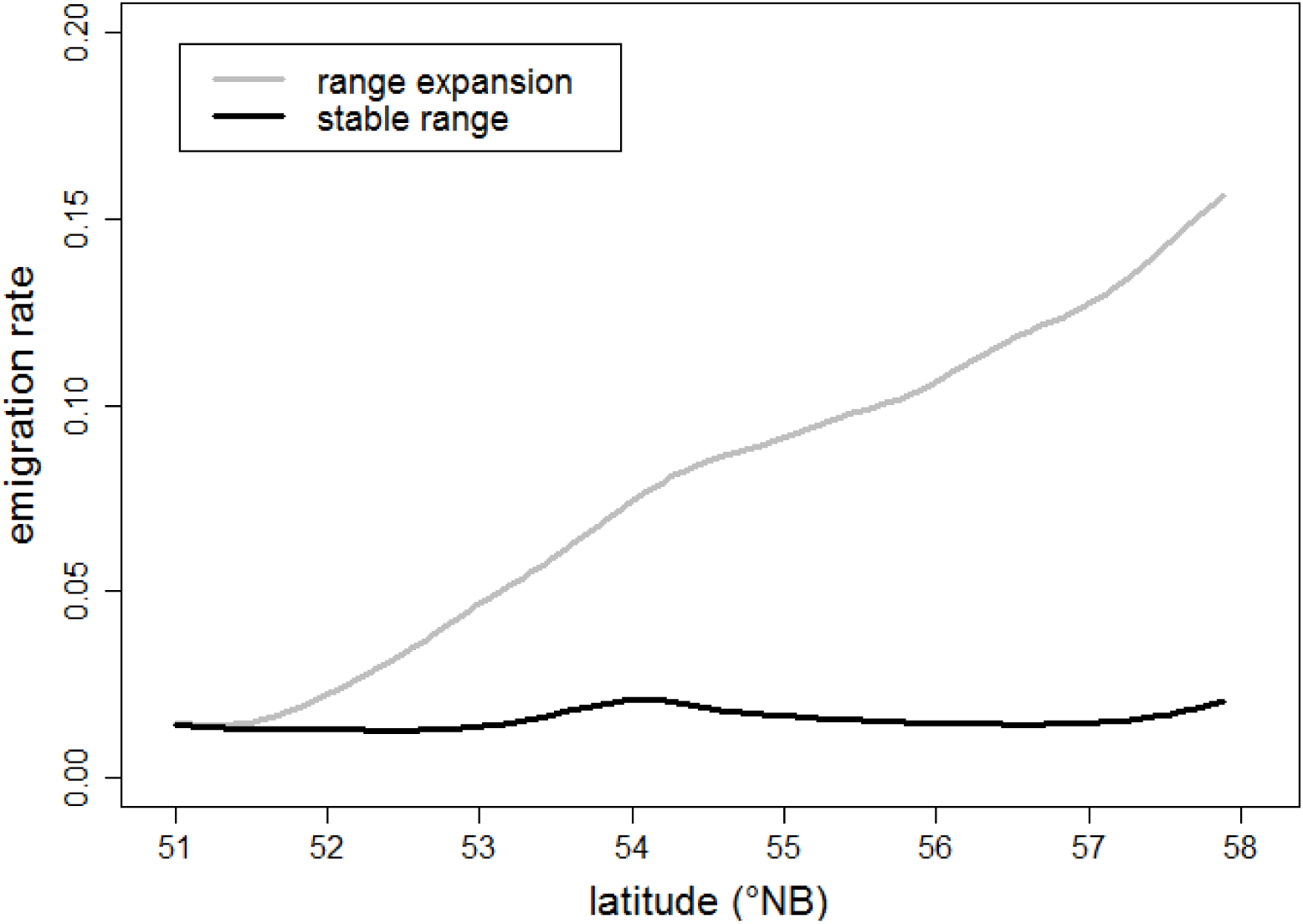
emigration rate as a function of latitude. The evolution of the emigration rate [embedded in the mites’ genome -see supplementary material S.3.0) is shown over the latitudinal range, for the model scenarios with range expansion *vs*. the model scenario with a stable range [here only for a trade-off setting of *max*. ±10% development, *max*. ±20% fecundity-see supplementary material S.3.5, figure S.3.5: d, h, 1). There is no latitudinal effect on emigration rate [*i.e*. dispersal) in the stable range scenario, while there is a clear increase in the range expansion scenarios.

**Figure S.6b:**
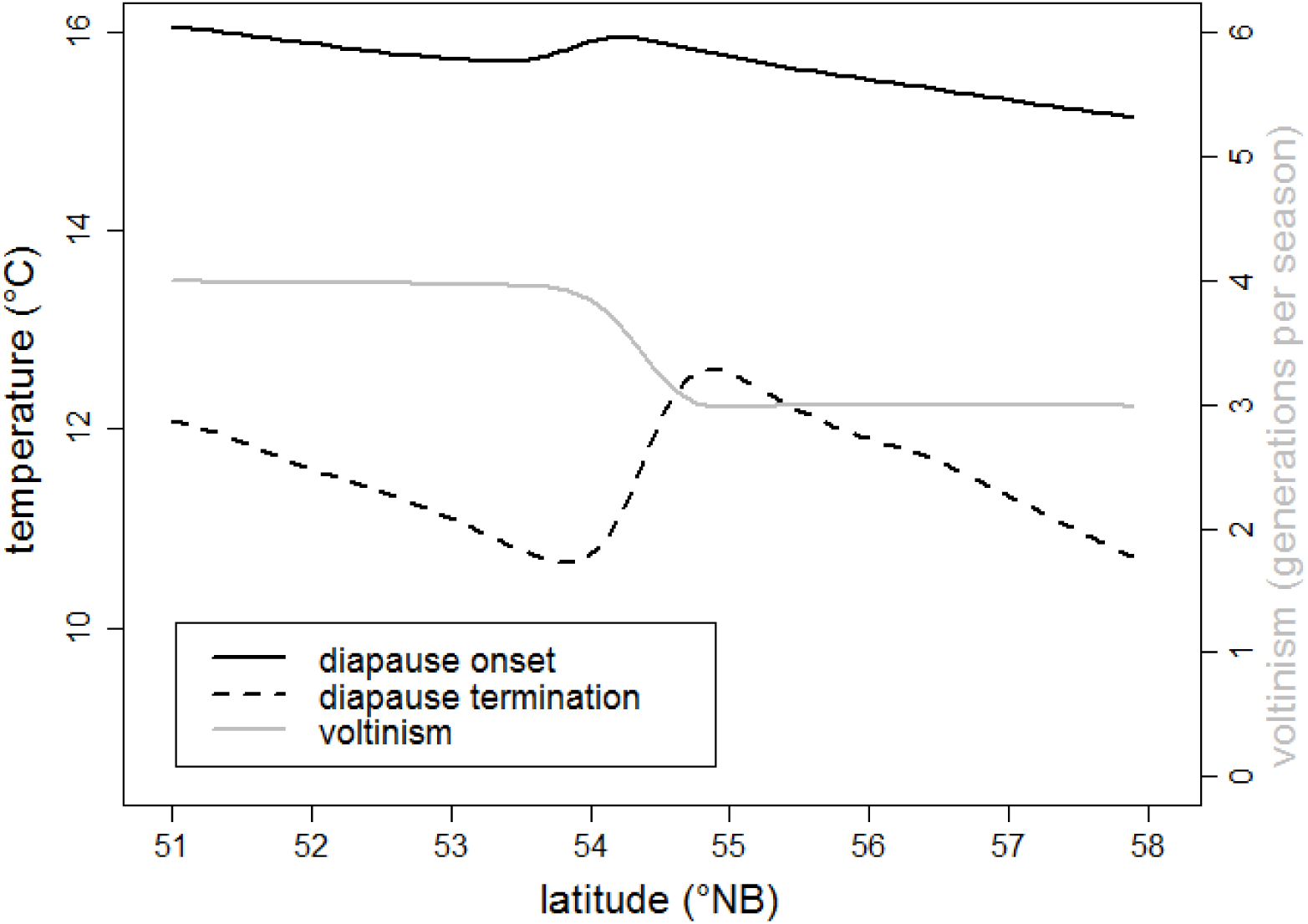
diapause behavior and voltinism as a function of latitude. Voltinism [grey line), the temperature of diapause onset [black line), and the temperature of diapause termination [dashed black line) are shown over the latitudinal range [here only for a stable range scenario with a trade-off setting of *max*. ±10% development *max*. ±20% fecundity - see supplementary material S.3.5, figure S.3.5: 1). When an environmental gradient is present, voltinism shows a stepwise decrease from south to north. Furthermore, the temperature of diapause termination is always lower than the temperature of diapause onset. Moreover, in those specific cases where a saw-tooth pattern in development time emerges [*e.g*. for a stable range scenario with a trade-off setting of *max*. ±10% development, *max*. ±20% fecundity, as shown here) there also emerges a saw-tooth pattern in the temperature of diapause termination, in synchrony with the step-wise decrease in voltinism.

### S.7: Trait correlations

**Table S.7:**
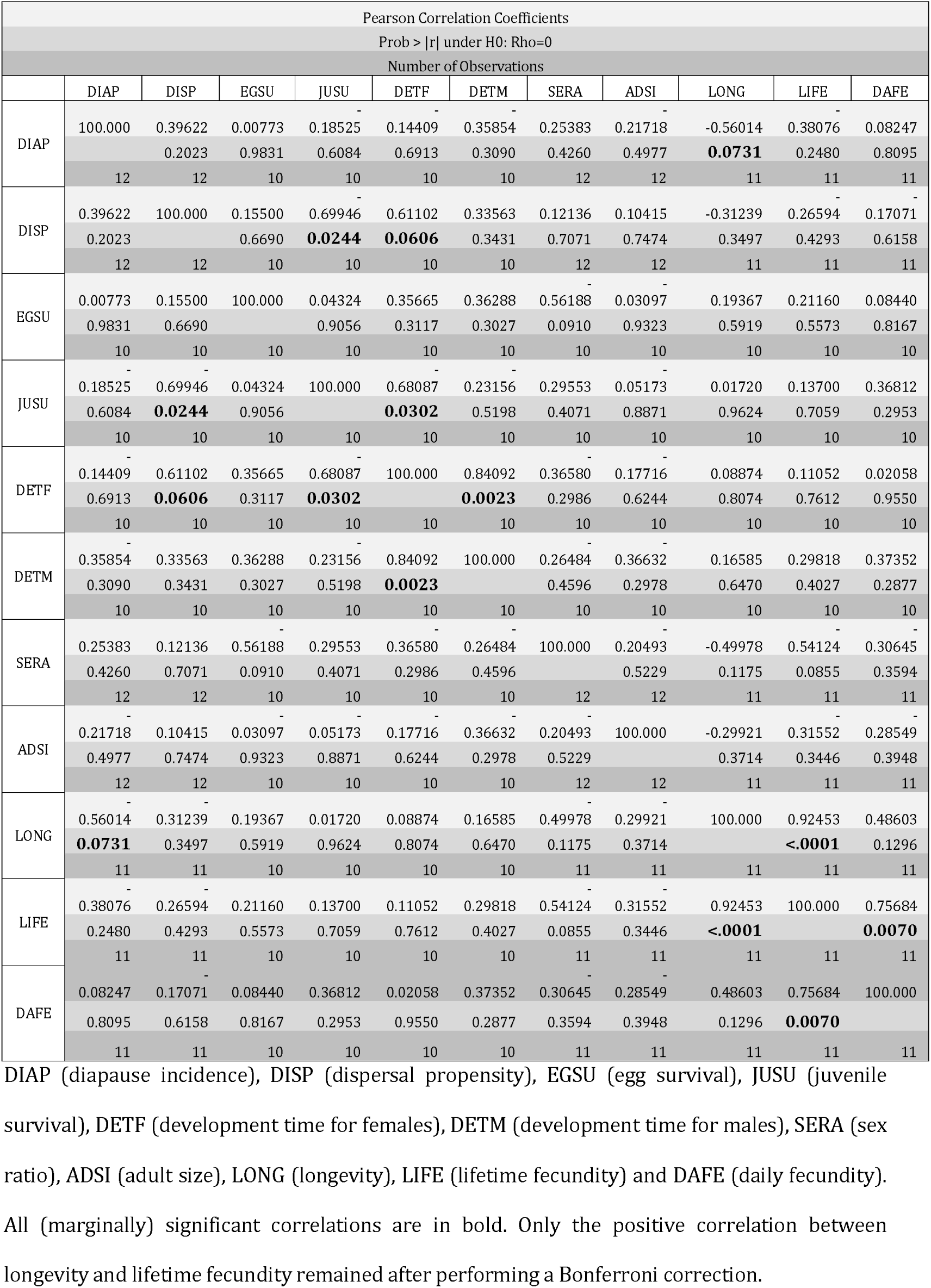
Pairwise correlations between all the life-history traits

